# Lysergic acid diethylamide pretreatment prolongs brain-stimulation induced neural activity changes

**DOI:** 10.64898/2025.12.16.694720

**Authors:** Lucas L. Dwiel, Mackenzi L. Prina, Elise M. Bragg, Matthew Company, Lucky L. Drucker, Laetitia R. Reduron, Bryan W. Luikart, Wilder T. Doucette

## Abstract

A leading theory for how psychedelics are able to produce robust clinical improvement and preclinical behavioral changes is that psychedelics act through neuroplastic mechanisms to induce lasting structural and functional alterations in neural circuits. However, psychedelics produce these effects across wide swaths of the brain. Based on our prior work, we hypothesized that engaging a specific brain circuit with focal brain stimulation during the window of enhanced plasticity produced by psychedelics would lead to more persistent alterations in the activity of that circuit. To test this, we administered either saline (SAL) or lysergic acid diethylamide (LSD) to rats 24 hours prior to electrical stimulation targeting the rat medial prefrontal cortex (mPFC). Brain activity was recorded before, during, and after stimulation using depth electrodes implanted in four bilateral frontostriatal regions. To assess changes in neural activity, we trained general logistic classifiers to distinguish between time points (e.g., pre-stimulation vs. post-stimulation) and then compared model performances across groups (e.g., LSD vs. SAL). As such, model performance represents the degree of difference between the two neural states (pre vs. post), and a significant difference between groups indicates a larger change in brain activity in one group. We found that LSD pretreatment, compared to SAL, resulted in larger and longer-lasting changes in brain activity following stimulation. Immunohistochemistry revealed that stimulation led to the activation of the mTOR signaling pathway, and the combination of LSD and stimulation led to alterations in perineuronal net (PNN) integrity. Regardless of pretreatment, brain activity states recorded during stimulation did not capture the brain state that persisted in the minutes or days that followed. This work has important implications for understanding the general effects of brain stimulation and provides strong support for the development of psychedelic-assisted brain stimulation approaches. By increasing the durability of brain stimulation-induced changes in activity, this approach could lead to a reduction in relapse rates, which currently limit the impact of non-invasive stimulation treatments.

## Introduction

Psychedelics have garnered considerable interest in the treatment of multiple psychiatric disorders with a small number of doses, sometimes a single dose, leading to significant alterations in mindset and behavior for months.[1–5] A leading hypothesis for this longevity is that psychedelics alter neural plasticity, allowing for long-lasting structural and functional changes in the brain. Many of the biomolecular pathways involved in psychedelic-induced neuroplasticity converge on mammalian target of rapamycin (mTOR), a kinase that regulates cell growth, protein synthesis, and transcription. Not only do psychedelics upregulate mTOR activation [6,7], but mTOR antagonists block the effects of psychedelics on dendritic growth [8,9], neural firing rate, and behavior [10]. Brain stimulation alone has also been shown to impact neuroplasticity through mTOR activation [11–13], making mTOR a possible convergence point between these interventions.

Despite robust acute effects, non-invasive brain stimulation outcomes typically do not last, with depression relapse rates as high as 80% within 6 months [14–17], 50% anxiety relapse within 3 months [18], and mixed effects in post-traumatic stress disorder (PTSD) [19,20]. To address this, stimulation protocols were lengthened, and/or patients would return to the clinic for additional sessions, creating additional burdens. Recent accelerated protocols, like the Stanford Neuromodulation Therapy for transcranial magnetic stimulation (TMS), have condensed multi-week stimulation sessions into 5 days[21], but only 47% of remitters maintain remission 12 weeks later[22]. There is also burgeoning evidence that pharmacological adjuvants targeting NMDA-dependent plasticity [23] enhance TMS response [24]. However, the persistence and replicability of these approaches remains to be established.

We have previously shown that infralimbic cortex (IL) brain stimulation 24 hours after LSD administration leads to larger and distinguishable changes in brain activity compared to stimulation following saline (SAL) [25]. The rodent IL target was selected due to its homology to the human subcallosal cingulate, an established stimulation target for depression. Based on these results and the known effects of psychedelics on neuroplasticity, we hypothesized that LSD would allow stimulation to produce longer-lasting changes in brain activity. We pretreated rats with either SAL or LSD, waited 24 hours, stimulated the IL, and quantified changes in local field potentials (LFPs) during and immediately after stimulation and across 7 days following stimulation. Given mTOR’s role as a potential convergence point and the role of perineuronal nets (PNNs) in governing neuroplasticity[26,27], we performed immunofluorescence staining to assess mTOR activation and PNN integrity.

## Methods

### Subjects and drugs

We used three cohorts of Sprague-Dawley rats (Charles River; Shrewsbury, MA) for these experiments (N = 38; 17 female, 21 male; 116-496 days old at the time of the experiment; see **Supplemental Information** for breakdown per cohort). We delivered 0.2 mg/kg LSD (Sigma Aldrich) or the equivalent volume sterile saline (SAL) via an intraperitoneal injection. We carried out all experiments in accordance with NIH Guide for the Care and Use of Laboratory Animals (NIH Publications no. 80–23) revised in 1996 and approved by the Institutional Animal Care and Use Committee (IACUC) at Dartmouth College.

### Electrode implantation

We anesthetized rats using an isoflurane/oxygen mixture (5% isoflurane for induction and 2% for maintenance) and stereotactically implanted them with custom electrode arrays[28] to record LFPs and deliver electrical stimulation. All electrodes targeted bilateral mPFC/infralimbic cortex (IL; AP 3.4 mm; ML ±0.75 mm; and DV -5.0 mm), orbitofrontal cortex (OFC; AP 3.4 mm; ML ±3.0 mm; and DV -6.0 mm), nucleus accumbens shell (NAcS; AP 1.2 mm; ML ±1.0; and DV -7.6 mm), and nucleus accumbens core (NAcC; AP 1.2 mm; ML ±2.4 mm; and DV -7.6 mm). All coordinates are relative to bregma.

### Stimulation

We used a Plexon stimulator (PlexStim, Plexon; Plano, TX) to deliver clinically based stimulation parameters (biphasic, 130 Hz, 90 μs pulse width, and ±200 μA) targeting bilateral IL. To obtain artifact-free LFPs during stimulation, every 50 seconds we turned the stimulation off for 10 seconds, repeating this either 60 or 120 times for 1 to 2 hours respectively; cohort 1 and 3 received 1 hour of stimulation, cohort 2 received 2 hours of stim. We verified current and voltage of stimulation before and after each session with a factory-calibrated oscilloscope (TPS2002C, Tektronix; Beaverton, OH). For sham stimulation, we plugged the rats in with the same stimulation cable, but did not pass current.

### Local field potential recording and processing

We recorded LFPs from freely-moving rats using a Plexon Omniplex digital acquisition system including commutators integrated into sound-proof enclosures (MedAssociates; Fairfax, VT) and simultaneous video acquisition using Plexon’s Cineplex system. We recorded LFPs at 20 kHz, downsampled to 2000 Hz, and removed 60 Hz noise with a notch filter. We quantified LFP features within 3-second windows (50% overlap), removing windows with artifacts (±2 mV) or active stimulation. We calculated power (*pwelch()*) and imaginary coherence (*cmtm()*)[29,30] across six frequency ranges: delta (Δ) = 1-4 Hz, theta (θ) = 5-10 Hz, alpha (α) = 11-14 Hz, beta (β) = 15-30 Hz, low gamma (lγ) = 45-65 Hz, and high gamma (hγ) = 70-90 Hz. From 8 recording locations, we obtained 48 power features and 168 coherence features (216 total).

### Experimental design

In cohorts 1 and 2, we injected each rat with SAL first, waited 24 hours, stimulated IL for 1 or 2 hours respectively, and collected LFPs during stimulation, for 1 hour after, and across 7 washout days after stimulation. After at least one week, we injected the same rats with LSD and repeated the protocol. We injected LSD second in all rats to avoid potential persisting effects of LSD+stim confounding the SAL control. In cohort 3, we split rats into 4 groups: SAL+sham (n = 5; 2 female, 3 male), SAL+sIL (n = 6; 3 female, 3 male), LSD+sham (n = 6; 2 female, 4 male), and LSD+sIL (n = 4; 3 female, 1 male). We followed the same protocol as before but euthanized the rats ∼20 minutes after 1 hour of stimulation for immunohistochemistry. Two rats, 1 male in SAL+sIL and 1 male in LSD+sham, did not contribute any neural data due to missing files. See **Fig. 1** for an experimental diagram of each cohort.

**Figure 1.**
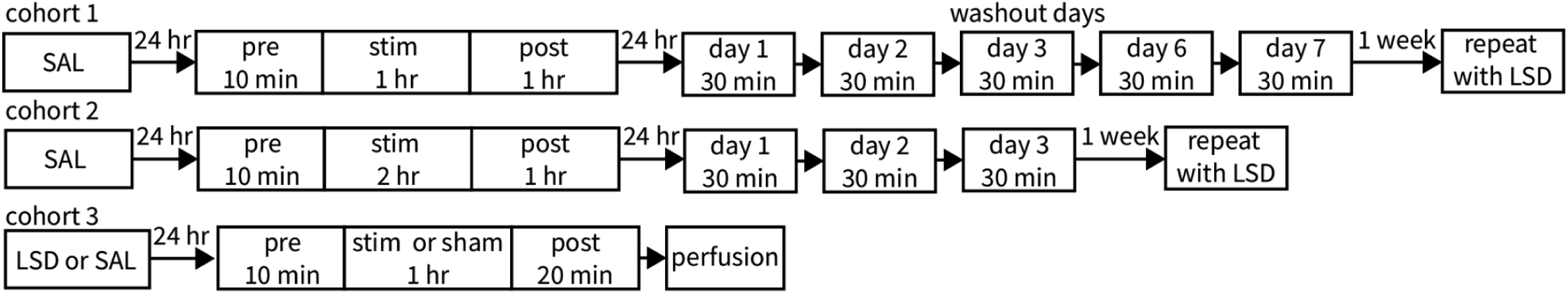
Experimental flowchart for each cohort.

### Model training and testing

To quantify the effect of LSD on brain stimulation, we trained logistic classifiers (*fitglm()*; Matlab 2025b; Natick, MA) to differentiate between: 1. pre- vs. post-stimulation LFPs (<u>pre vs. post</u> models), 2. pre-stimulation vs. during-stimulation LFPs (<u>pre vs. stim</u> models), and 3. SAL vs. LSD post-stimulation or during-stimulation LFPs (<u>SAL vs. LSD</u> models). For each model, we balanced data such that each rat contributed equally, split 80/20 for training/testing, and quantified performance as area under the receiver operator characteristic curve (AUC; *perfcurve()*). We repeated all modeling 100 times to account for subsampling variability. To assess persistence, we tested pre vs. post and pre vs. stim models on washout day data. To assess similarity between LSD and SAL brain states, we cross applied models. We characterized model variability across time and rats by testing models on 5 minute bins of data per rat. For single feature analysis, we built and tested individual models per feature. We built sex-specific models to test for sex differences.

To establish null distributions, we built permuted models with shuffled group labels. As an additional control, we obtained sham stimulation recordings from a subset of rats (n = 14; 7 female and 7 male) before any injection, built analogous models, and tested on subsequent days. The washout data (24, 48, and 96 hours) did not perfectly match the data in the LSD or SAL groups (24, 48, 72, 144, and 196 hours) so we averaged performances across all washout days to obtain an estimate of SHAM model performance (AUC = 0.67±0.05; **Supplementary Figure 1**). The true null likely falls between the permuted and sham estimates.

### Statistical analysis

We compared model performance distributions using permutation tests, p = (1+m)/(1+b), where m is the number of times that the permuted distribution outperformed the mean of the real distribution and b is the number of values in the permuted distribution. [31] We used linear mixed effects models (AUC∼Group*Time) to compare model performances across washout days, with post-hoc permutation tests. We compared variability of model performance (standard deviation of AUCS across time and rats) using two-sample t-tests. For washout feature analysis, we identified features exceeding the 95th percentile of permuted performance and compared these between LSD and SAL groups. We used Pearson correlations (*corrcoef()*) to assess whether the same features differentiated both stim and washout conditions.

### Feature visualization

To visualize individual LFP features across time, we z-scored feature values to their baseline levels and then averaged across rats either in 5 minute intervals for during and post-stimulation or across the 30 minute washout day recording.

### Velocity

We tracked rat movement with DeepLabCut[32], computed a centroid from four body landmarks, and calculated velocity (pixels/minute). Be built pre vs post models of velocity following the same procedure as LFPs to determine whether velocity changes alone could explain model performance.

### mTOR/PNN

#### Immunohistochemistry and image analysis

After collecting ∼20 minutes of post-stimulation LFPs, we sacrificed and perfused the rats with PBS and 4% paraformaldehyde. We extracted the brains and stored them in 4% paraformaldehyde. After slicing and staining, we imaged all sections with a Zeiss LSM800 laser-scanning confocal microscope, and analyzed fluorescence intensity with ImageJ/Fiji (see **Supplemental Information** for full staining and imaging protocols). We assessed group differences with linear mixed effects models (Intensity∼Sex+Age+Group) per brain region, with *coefTest()* for post-hoc comparisons.

## Results

### LSD allows for a more stable and longer-lasting stimulation-induced brain state

Models built and tested on LFPs from before and 1 hour after stimulation (pre vs. post) outperformed permuted models in both groups (p < 0.01; **Fig. 2A**) with no effect of sex (**Supplementary Figure 2**). The post-stimulation brain activity of the LSD group was significantly easier to differentiate from pre-stim than that of the SAL group (p < 0.01; **Fig. 2A**). The change in brain activity detected by these models persisted across washout days in both groups compared to permuted models (**Supplemental Information**) but only the LSD group outperformed the overfit control models (**Fig. 2B**). Comparing LSD to SAL revealed that LSD significantly increased model performance (β = 0.12 [0.03 0.22], SE = 0.028, t(6) = 3.16, p = 0.020) with no effect of time or interaction between group and time (**Fig. 2B**). Post-hoc permutation tests showed the group difference occurred over the first 3 days (p = 0.0099, 0.030, and 0.020 respectively). The fact that the SAL data overlaps with the sham model performance (**Fig. 2B**; heavy black line) suggests that there are no persisting effects of stimulation alone. Although there is a decrease in velocity from pre to post stimulation, this difference does not allow a model built on velocity data to outperform chance in differentiating pre from post or pre from washout data (**Supplementary Figure 3**). Therefore, the performance of the pre vs. post models on both post and washout data can not be entirely explained by changes in velocity.

**Figure 2.**
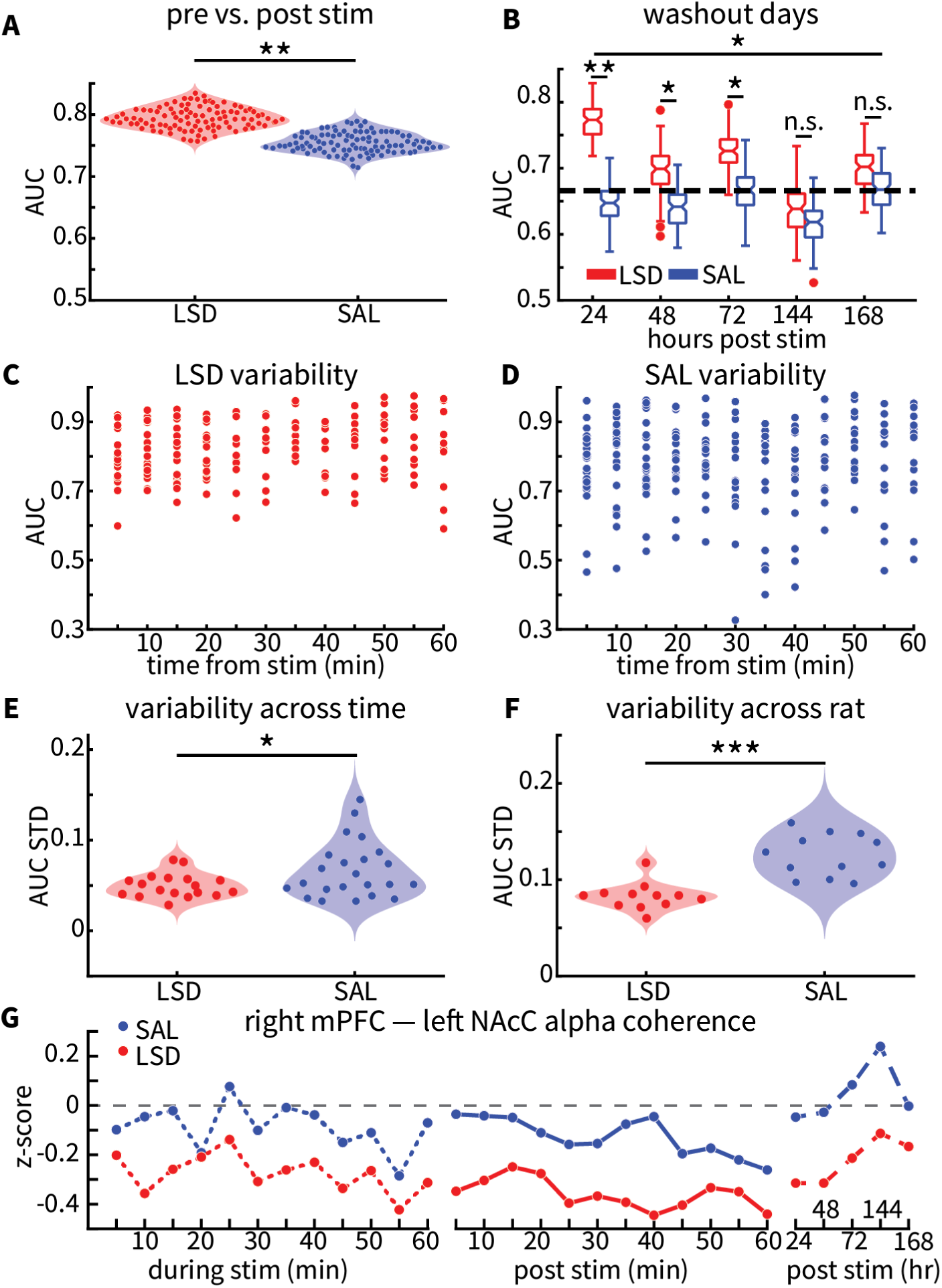
LSD pretreatment allows for a large, longer-lasting, and stabler stimulation-induced brain state. **A.** Models built to differentiate pre-stimulation from post-stimulation neural data outperform chance (not shown) in either the LSD group (red) or saline group (SAL; blue) and there is a larger difference between pre and post data in the LSD group compared to the SAL group. **B.** LSD models outperform SAL models on washout days, for up to 3 days after stimulation. Heavy dashed black line indicates the estimated performance of overfit models from **Supplementary Figure 1**. **C-F.** The performance of the models in A on individual rats outperforms chance in LSD (**C**) and SAL (**D**) with less variability (standard deviation) in performance across time (**E**) and rats (**F**) in LSD vs. SAL for the first hour after stimulation. **G.** A representative feature, z-scored from baseline levels, shows shifts during stimulation in the LSD group that remain for 60 minutes and multiple days after stimulation. *** p < 0.001, ** p < 0.01, * p < 0.05, and n.s. = not significant.

Not only was brain activity more changed in the LSD group for the first hour after stimulation, but the performance of the models on individual rats was less variable when given LSD (**Fig. 2C-D**). Using two-sample t-tests shows a decreased standard deviation in AUC for each rat across time (mean difference = -0.017 [-0.033 -0.0013], t(39) = -2.2, p = 0.034; **Fig. 2E**) and for each 5 minute time interval across rats (mean difference = -0.042 [-0.058 -0.027], t(22) = -5.6, p < 0.001; **Fig. 2F**). A representative feature (right mPFC to left NAcC alpha coherence) in the pre vs. post models for the LSD group illustrates the shift persisting for days in the LSD group while returning to baseline immediately in the SAL group (**Fig. 2G**).

### LSD interacts with stimulation to produce a similar, but distinct brain state

To determine if the same features were persistently changed regardless of pretreatment, we compared single feature pre vs. post model performance on washout data. Of 14 LSD features exceeding the 95th percentile of permuted performance, 12 also outperformed the corresponding SAL feature (**Fig. 3A).** These features were predominantly corticostriatal coherence and accumbal power (**Fig. 3B**) that persisted into the washout days. Comparing the performance of these single feature models between the original pre vs. post data and the washout data from 24 hours after stimulation revealed that all ten models substantially increased in performance in the LSD data compared to the SAL data (**Fig. 3C**), indicating that these features were not only changed by stimulation for the first hour, but continued to change in the same direction over the next 24 hours–moving further from baseline. Cross-applying models between groups decreased performance, though not to chance levels, indicating that LSD alters the effects of stimulation while retaining some overlap between groups(**Fig. 3D**). Furthermore, we built models to directly differentiate LSD from SAL during and after stimulation also outperformed chance (**Fig. 3E**).

**Figure 3.**
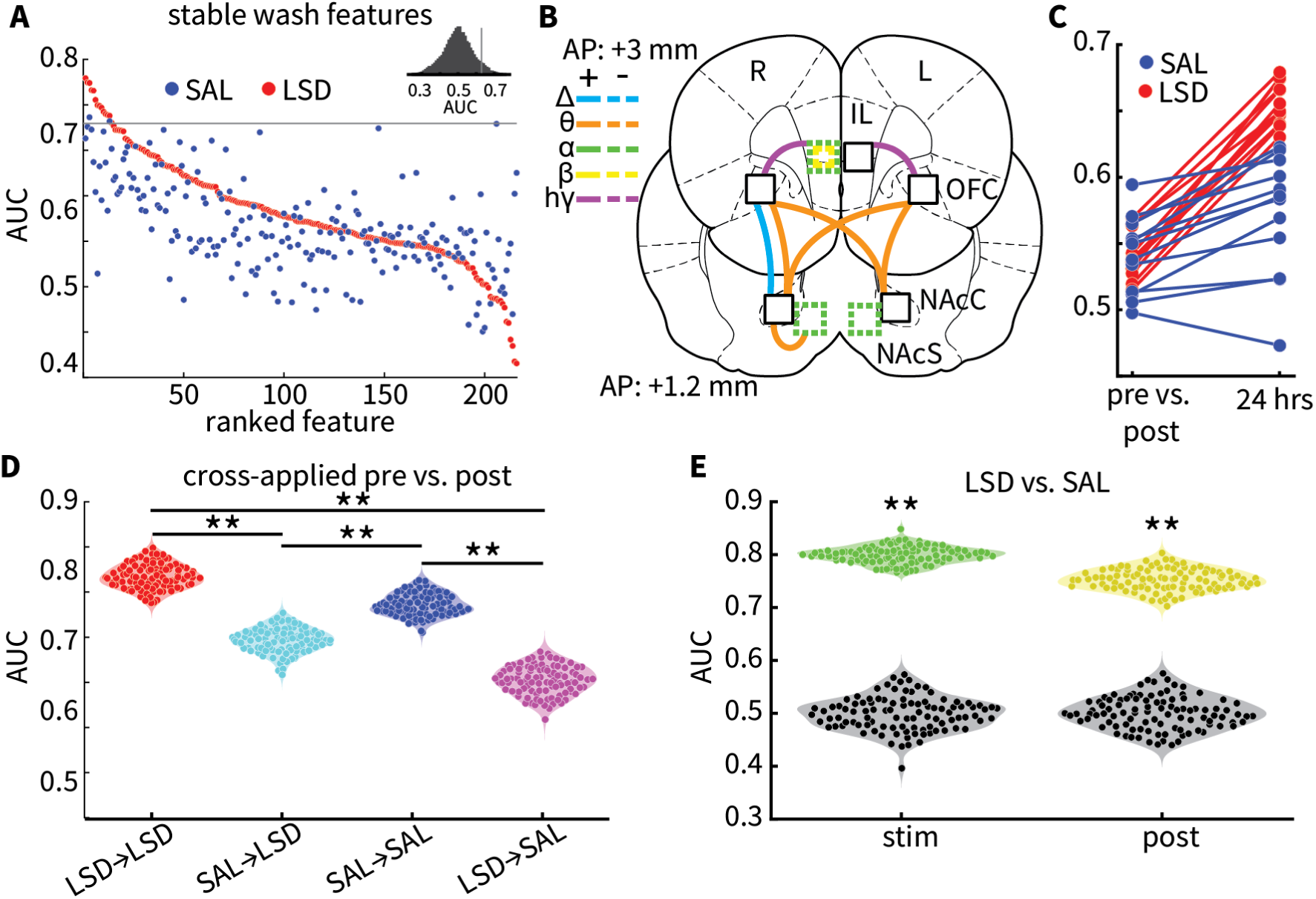
LSD alters the effects of stimulation. **A.** Performance of single feature pre vs. post models on washout days, sorted by performance on LSD data. Red dots come from LSD data and blue from SAL. Horizontal grey line indicates 95% threshold of performance based on the distribution of LSD permuted single feature models (top inset). **B.** Map of the top 12 LSD features from **A** that outperform corresponding SAL models. Solid vs. dashed lines indicate if a given feature is increased or decreased post stimulation compared to pre stimulation. Lines connecting brain regions indicate coherence between those regions and colored outlines indicate power in that region. Color indicates frequency range. **C.** Comparison of the change in single feature model performance from the first hour after stimulation to 24 hours after for the top 12 features from **A** and **B**. LSD data in red and SAL in blue. **D.** Cross-applying models lowers performance in the first hour after stimulation compared to training and testing models within datasets. **E.** Models built to differentiate between LSD and SAL changes during stim (green) and during the hour after stimulation (yellow) outperform chance. ** p < 0.01.

### Brain states induced by stimulation evolve over time

We were also able to build models to differentiate between pre-stim LFPs and LFPs during stimulation (pre vs. stim) that not only outperformed permuted models (p < 0.01; **Fig. 4A**), but also their corresponding pre vs. post models for both LSD (p < 0.01; **Fig. 4A**) and SAL groups (p < 0.01; **Fig. 4A**). However, applying these pre vs. stim models to pre vs. post data significantly decreased performance (p < 0.01; **Fig. 4A**). Further, we compared the performance of the pre vs. post and pre vs. stim models on washout days and found that the pre vs. stim models performed worse than pre vs. post models in the LSD group (β = -0.13 [-0.22 -0.042], SE = 0.037, t(6) = -3.60, p = 0.011; **Fig. 4B**), but not the SAL group. A representative feature, left NAcS to right NAcS delta coherence, shows a robust decrease in both groups while stimulation is active that immediately returns to baseline levels when stimulation is turned off and throughout the washout days (**Fig. 4C**). Last, comparing the performance of single feature models from the LSD pre vs. stim models on pre vs. stim data and pre vs. wash data shows no significant correlation indicating that a feature that can detect when stimulation is on does not tend to also predict persistent changes and vice versa (**Fig. 4D**).

**Figure 4.**
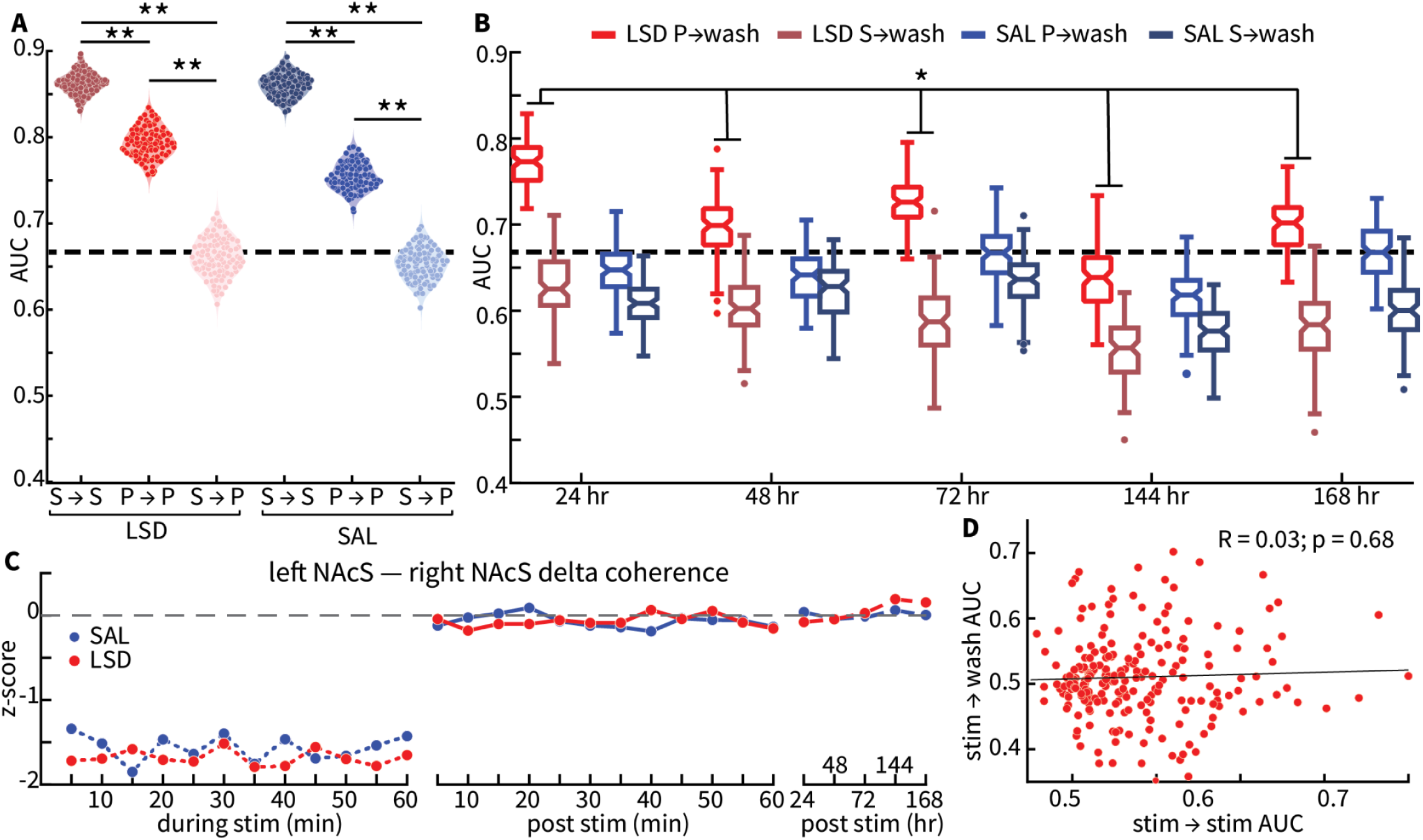
Brain states induced during stim do not persist, but states that emerge immediately after stim do. **A.** Building and training models within stim or post (S → S; P → P) outperforms a model trained on stim and tested on post (S → P). All models outperform chance. **B.** Models built on post data from the LSD group outperform models built on stim data from the LSD group. **C.** Representative features, z-scored from baseline levels, showing the dynamics of a feature changed by stimulation. Both groups show a decrease in left nucleus accumbens shell (NAcS) — right NAcS delta (Δ) coherence during stimulation, that immediately rebounds to baseline levels after stimulation and remains there through washout days. **D.** There is no correlation between single feature model performance (AUC) of models trained on pre vs. stim data and tested on pre vs. stim (x axis) or pre vs. wash (y axis) for LSD. ** p < 0.01, * p < 0.05, and n.s. = not significant. Heavy dashed black line in A and B indicates the estimated performance of overfit models from **Supplementary Figure 1**.

### Cohort comparison

Since the two cohorts had different amounts of stimulation (1 hour for cohort 1 and 2 hours for cohort 2), we replicated all of the above analyses for each cohort on its own and despite an additional hour of stimulation in cohort 2, model performance did not differ between cohorts for either acute or washout comparisons (**Supplementary Figure 4**).

### mTOR activation and PNN integrity

We found group differences in mTOR activation (pS6 intensity) in IL (F(3,1156) = 4.319, p<0.005; **Fig. 5A**) and mPFC (F(3, 749) = 6.403, p < 0.001; **Fig. 5B**), but not DG (F(3,260) = 1.422, p = 0.24; **Fig. 5C**). We found an effect of age in all regions, with increased mTOR activation in older rats: IL (F(1,1156) = 9.026, p < 0.003), mPFC (F(1,749) = 35.694, p <0.001), and DG (F(1,260) = 19.011, p < 0.001). We also found an effect of sex in IL (F(1,749) = 9.935, p < 0.002), with more activation in males. Post-hoc analysis revealed that in both IL and PL, mTOR activation was highest in LSD-sIL and driven primarily by stimulation (**Fig. 5A-C** and **Supplemental Information** for p-values). However, the absence of a SAL-sIL increase over SAL-sham suggests an interaction between LSD pretreatment and stimulation.

**Figure 5.**
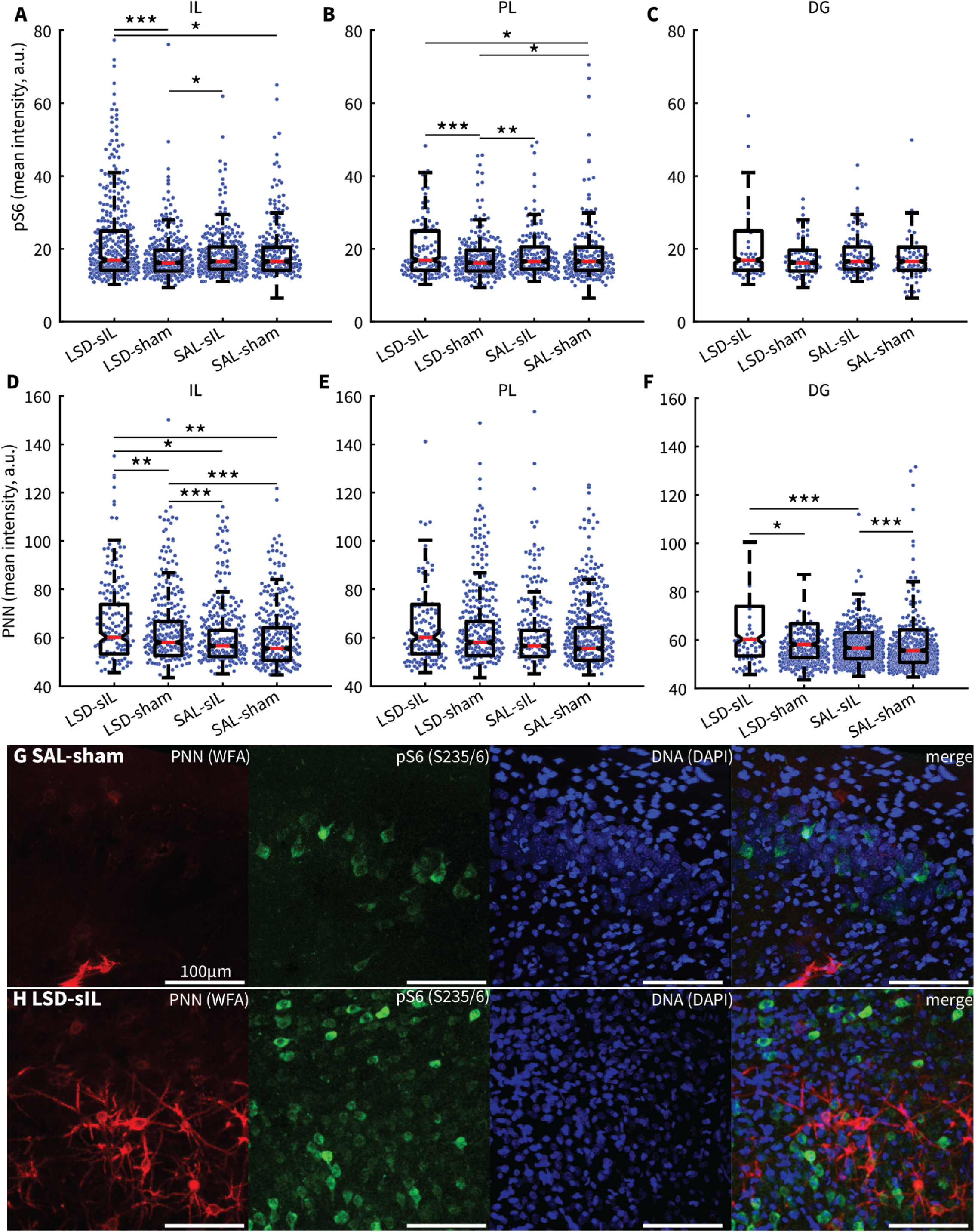
LSD-sIL elevates mammalian target of rapamycin (mTOR) and perineuronal nets (PNN) in infralimbic cortex (IL). **A-C.** Phospho S6 staining intensity as a marker of mTOR activation across the 4 groups in 3 brain regions: IL (**A**), prelimbic cortex (PL; **B**), and dentate gyrus (DG; **C**). **D-F.** PNN staining across the 4 groups in the same 3 brain regions as **A-C**. **G-H.** Representative images of staining from a rat in the SAL-sham (**G**) and LSD-sIL (**H**) groups. Wisteria floribunda lectin (WFA) staining as a marker for PNN integrity in red, S235/6 staining for pS6 in green, DAPI for DNA in blue. Scale bar is 100 μm. *** p < 0.001, ** p <0.01, * p < 0.05

We found group differences in PNN status (WFA intensity) in IL (F(3,792) = 9.968, p <0.001; **Fig. 5D**) and DG (F(3,1519) = 8.971, p <0.001; **Fig. 5F**), but not PL (F(3,784) = 1.287, p = 0.28; **Fig. 5E**). We found an effect of sex in all regions with more PNNs found in males: IL (F(1,792) = 8.312, p = 0.004), PL (F(1,784) = 6.63, p = 0.01), and DG (F(1,1519) = 7.192, p = 0.007). Last, we found an effect of age in PL (F(1,784) = 13.598, p < 0.001) and DG (F(1,1519) = 5.461, p = 0.02) with increased PNNs in older rats. Post-hoc analysis revealed that in IL, LSD-sIL had greater PNN intensity than all other groups, and LSD-sham exceeded both SAL groups (**Fig 5D-F** and **Supplemental Information**). In DG, both stimulation groups exceeded their sham counterparts. Thus, LSD alone increases PNN integrity in IL, and interacts with stimulation to produce an even larger increase.

## Discussion

Here we show that LSD pretreatment allows for brain stimulation to push the brain further away from baseline, and that this altered state persists for days longer than after SAL. This means that psychedelics may allow for brain stimulation therapies like TMS to have longer lasting effects, reducing the high relapse rates currently limiting these non-invasive stimulation approaches. We also show that the brain state during stimulation differs from the state that evolves after stimulation ends; importantly, it is the post-stimulation state that persists for days and would support any therapeutic effect. This means that biomarkers recorded during stimulation would poorly reflect lasting changes.

### LSD allows for larger and longer-lasting effects of brain stimulation

By stimulating the IL 24 hours after LSD, we produced stable, long-lasting changes in brain activity distinguishable from those following SAL. The LSD pre vs. post model had higher and more stable performance both in the hour after stimulation and across subsequent days. Moreover, although there is overlap in the effects of stimulation regardless of pretreatment, LSD also allows for stimulation to produce novel effects on brain activity. These results suggest that psychedelics could serve as adjuvant therapies alongside neuromodulation to promote lasting changes in brain activity and mood. Although we included a sham model to capture non-intervention-related LFP changes across time, we did not collect washout LFPs from rats given only LSD. Therefore, we can not be certain that persistent changes are not at least partly due to residual LSD effects nor stacked stimulation sessions. However, we previously showed that LSD’s acute effects on LFPs are largely resolved within 24 hours[33], and since SAL+sIL does not produce persistent changes, it seems unlikely that the first stimulation session would be interacting with the LSD+sIL session 1 week later to produce the observed persisting effects. However, it could be the case that the first stimulation session is producing latent changes in the brain or persisting effects that are not captured by LFP power and coherence. Future work using counterbalanced designs or fully between-subjects groups will help address this limitation.

We found at least a 3-fold increase in the duration of stimulation-induced brain state changes with LSD pretreatment (i.e., SAL+stim altered brian activity for at least 1 hour, while LSD+stim produced changes for at least 3 days). If translatable, this could extend therapeutic effects for additional months to years. Granted, the differences between our single session protocol in wild-type rats and multi-session clinical courses in patients warrant caution in direct translation. [34–37]

Others have demonstrated the potential for neuroplastic modulators, (e.g., D-cycloserine) to enhance the effects of TMS [38,39] and other serotonergic psychedelics, like 2,5-dimethoxy-4-iodoamphetamine (DOI), have been shown to enhance/modulate long-term potentiation (LTP) [40–42]. Interestingly, when ketamine or psilocybin have been paired acutely with TMS pulses, neither showed impact on evoked response complexity. [43,44], suggesting that concurrent protocols may not allow sufficient time for engaging neuroplastic processes. We hypothesize that psychedelic pretreatment produces a malleable state in which dendritic spines are more likely to be ‘immature’ and neurons primed for neuroplasticity. By applying stimulation in this state, more robust changes in brain circuits are produced.[45] The timing between psychedelic and stimulation is likely a critical factor: psychedelics alter spine maturity within 24 hours [46,47], but this shift disappears by 34 days [48,49].

The neural features that remain altered for days are predominately coherence measures, suggesting that stimulation after LSD produces lasting changes in connectivity between brain regions—particularly increased OFC-NAc connectivity despite IL being the stimulation target. Future work should test whether stimulating different frontostriatal nodes produces the same effects, whether persistence depends on stimulation targeting or parameters (i.e., frequency and duration), and whether shorter-acting psychedelics like psilocybin or DMT produce comparable results.

### Brain states induced by stimulation evolve over time

The brain state during stimulation is not the same as the state arising in the hours or days after. At the individual feature level, features most changed during stimulation are rarely the same as those changed afterward. Instead, we find features with marginal changes during stimulation that change when stimulation is turned off and remain changed for days. Clinically, deep brain stimulation (DBS) effects on brain activity evolve through time, with features predictive of clinical outcome acutely decreasing with stimulation followed by a gradual increase that correlates with clinical benefit [50]. Similarly, in TMS for cocaine use disorder, lasting shifts in brain state dynamics correlated with a sustained decrease in cravings [51]. This implies that optimizing stimulation parameters can not rely solely on acute biomarkers; effective optimization may require multiple sessions over days or weeks.

### Clinical outcomes and brain state stability

The theoretical framework of this work is that successful psychiatric treatment depends on lasting shifts in brain state, and relapse occurs when the brain returns to its pre-treatment state. This is supported by evidence that DBS induced changes in brain activity correlate with clinical benefit for depression [52], that depression relapse correlates with decreased network resilience/stability,[53,54] and that residual symptoms in remitted depression correlate with maintained connectivity disruptions [55]. However, the relationship between healthy, remitted, and relapsed brain states is complex as relapse may reflect an inability to establish stable network structure rather than a return to a specific state.[56] Our work suggests that psychedelics can help stimulation produce such stable changes.

### mTOR is upregulated by LSD and stimulation

Stimulation led to greater mTOR activation than control conditions, and LSD pretreatment produced the highest levels. Although LSD-sIL did not significantly exceed SAL-sIL in mTOR activation, the absence of an expected SAL-sIL increase over SAL-sham suggests an interaction between LSD and stimulation, even 24 hours after drug administration [57]. This is consistent with a growing literature on psychedelic modulation of mTOR [58–62] and with the established role of mTOR in the neuroplastic and antidepressant effects of ketamine [63–74] and other antidepressants [75–80]. Prior work has shown that one hour of IL stimulation upregulates AKT and CREB phosphorylation through mTOR [81], and that chronic stimulation protocols similarly engage mTOR signalling alongside behavioral improvement [82,83]. However, mTOR is but one node in a complex network [84,85], with BDNF/TrkB as key upstream regulators [86,87]. Both stimulation [88–93] and psychedelics [57,94] engage BDNF/TrkB signalling, suggesting multiple convergent pathways. Our findings of increased mTOR activation in males and older rats is consistent with known effects of sex and age on mTOR signalling. [95–101]

### LSD and stimulation interact to increase PNN integrity

LSD alone increased PNN staining intensity in IL, and LSD pretreatment interacted with stimulation to produce an even larger increase. PNNs regulate synaptic transmission [102,103] and plasticity, with their accumulation signalling closure of critical periods [104,105], and they have been implicated in early life adversity [106,107], memory [108,109], and neuroinflammation [110]. Although little is known about psychedelic effects on PNNs, chronic ketamine has been shown to both decrease [111] and increase [112] mPFC PNNs with increased PNNs showing altered morphology. We found that although LSD-sIL increased PNNs in DG compared to LSD-sham and SAL-sIL, this effect was not greater than SAL-sham, suggesting hippocampal effects may be more stimulation-dependent. A more thorough examination of PNN dynamics under psychedelic treatment would help determine whether psychedelics first disrupt PNNS to enable plasticity and then facilitate their reassociation, thereby stabilizing synaptic changes. Our findings of increased PNNs in males and older rats are consistent with, though not easily disentangled from, the complex and sometimes contradictory literature on age and sex effects. [113–124]

## Conclusion

We have confirmed that LSD given 24 hours before brain stimulation allows stimulation to induce larger changes in brain activity that persist for days rather than hours. Further, LSD and brain stimulation may interact through upregulation of mTOR signaling and increased PNNs. Combined with the known effects of psychedelics on neuroplasticity, these findings suggest that psychedelics produce a state of enhanced neuroplasticity that can be acted upon by targeted brain stimulation to create lasting changes across broader neural circuits—potentially enabling the persistent shifts in mindset and behavior sought in neuropsychiatric treatments.

## CRediT

Conceptualization: LD, WD, BL

Methodology: LD, WD, BL

Software: LD

Formal Analysis: LD, MC

Investigation: LD, MP, L Drucker, LR

Resources: WD, BL

Data Curation: LD

Writing - Original Draft: LD

Writing - Review & Editing: LD, WD, MP, LR, MC

Visualization: LD, MP

Supervision: LD, WD, BL

Project Administration: LD

Funding Acquisition: LD

---

**Supplementary Figure 1.**
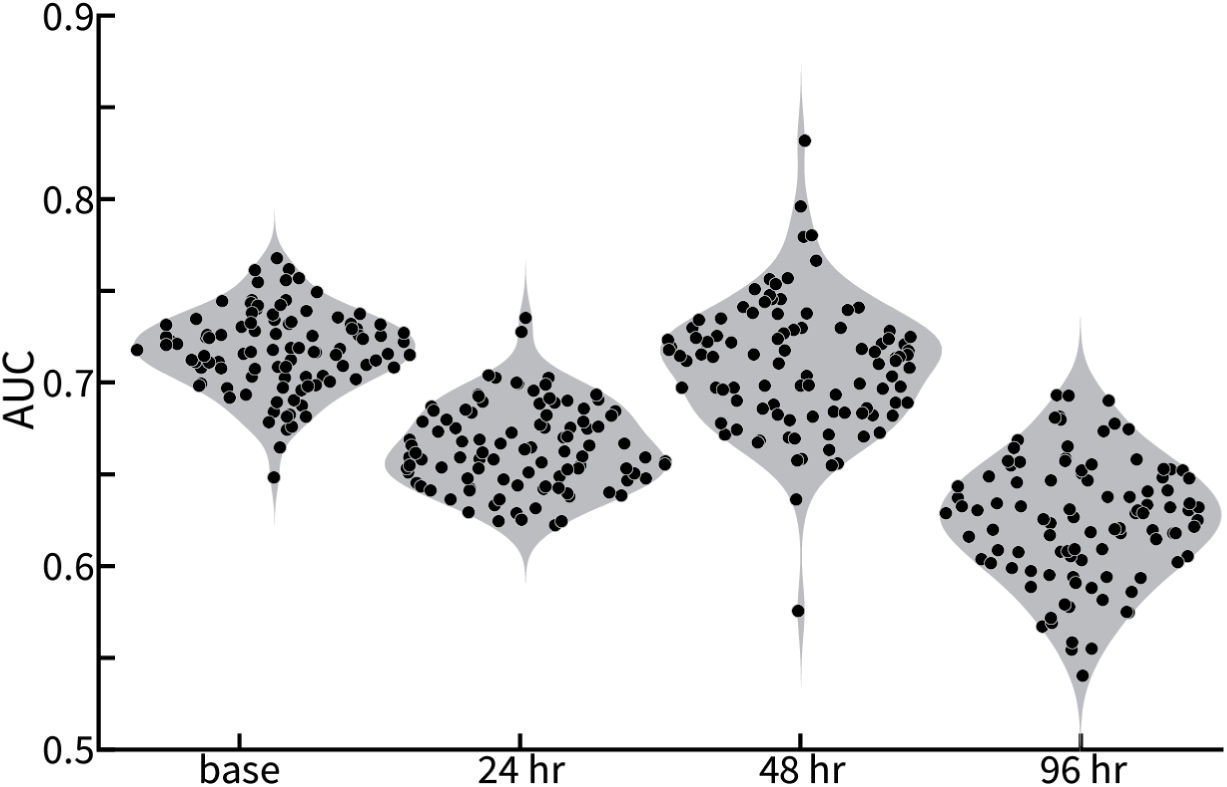
Sham stimulation models built from recordings before any injections showing performance across time. We averaged performances on 24, 48, and 96 hours after the baseline data to provide an estimate of model performance on washout days due to non-interventional changes through time.

**Supplemental Figure 2.**
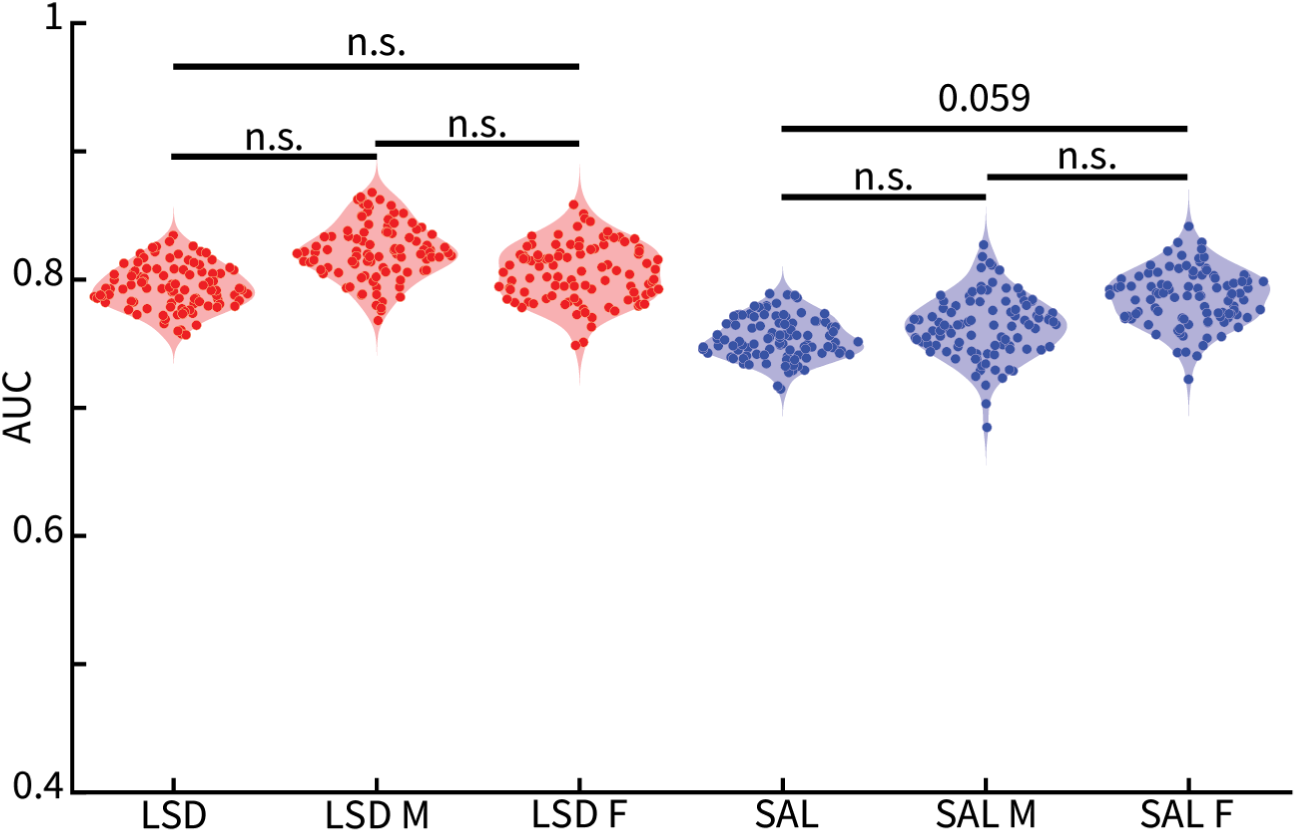
Pre vs. post models generalize across sex. Testing the pre vs. post models built on data from both sexes (Fig. 1A) on each sex separately revealed no difference in performance between sexes nor between each sex and the full test set.

**Supplemental Figure 3.**
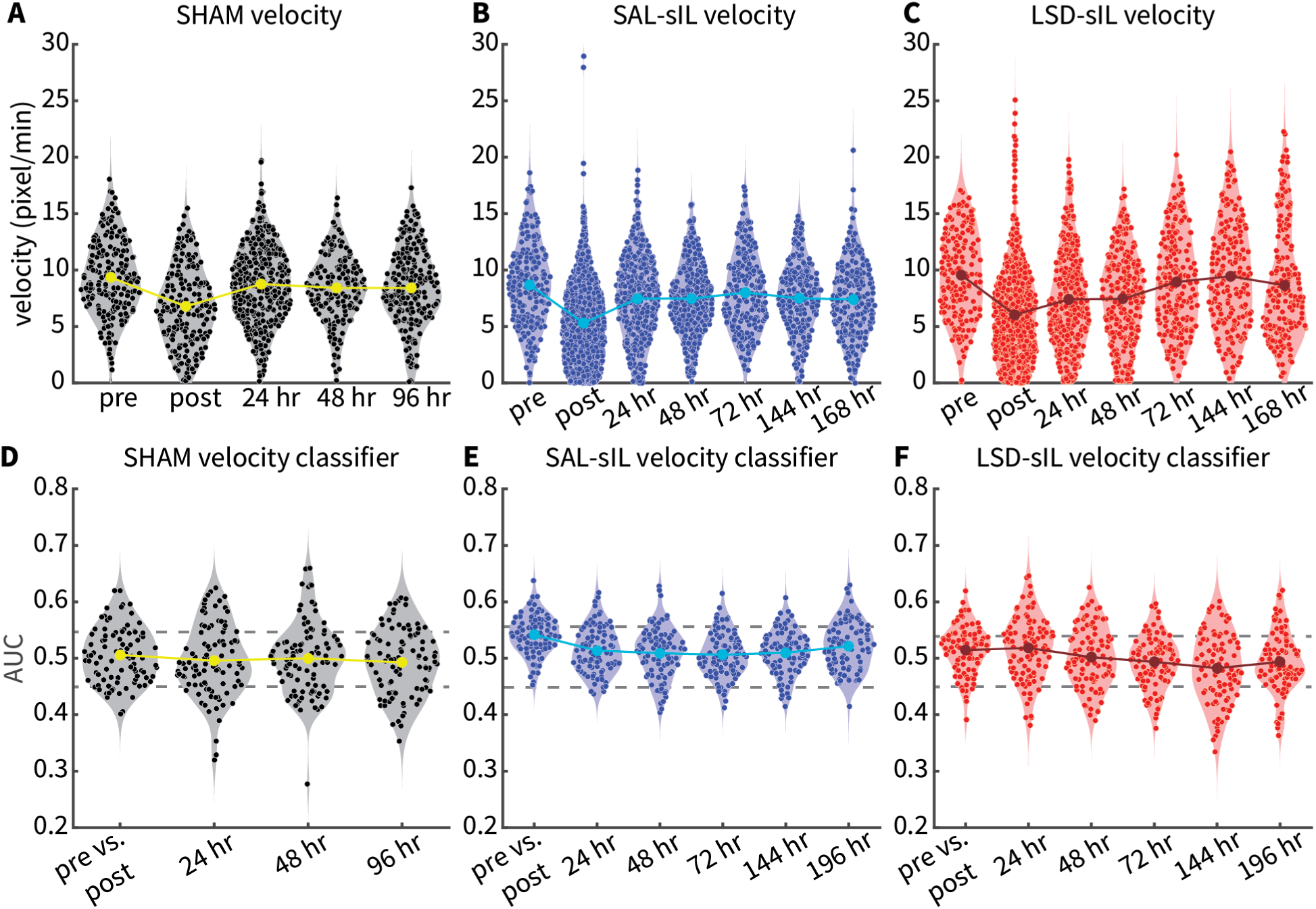
Velocity does not significantly contribute to model performance. **A-C.** Velocities (pixel/min) for each group across time corresponding to intervals used for LFP-based classifiers. Large circles and solid lines indicate average velocities. **D-F.** Performances of classifiers built from velocity data then tested across time. Large circles and solid lines indicate average AUCs and dashed grey lines indicate the mean±standard deviation of permuted classifier performance. No models outperform their corresponding permuted range.

**Supplemental Figure 4.**
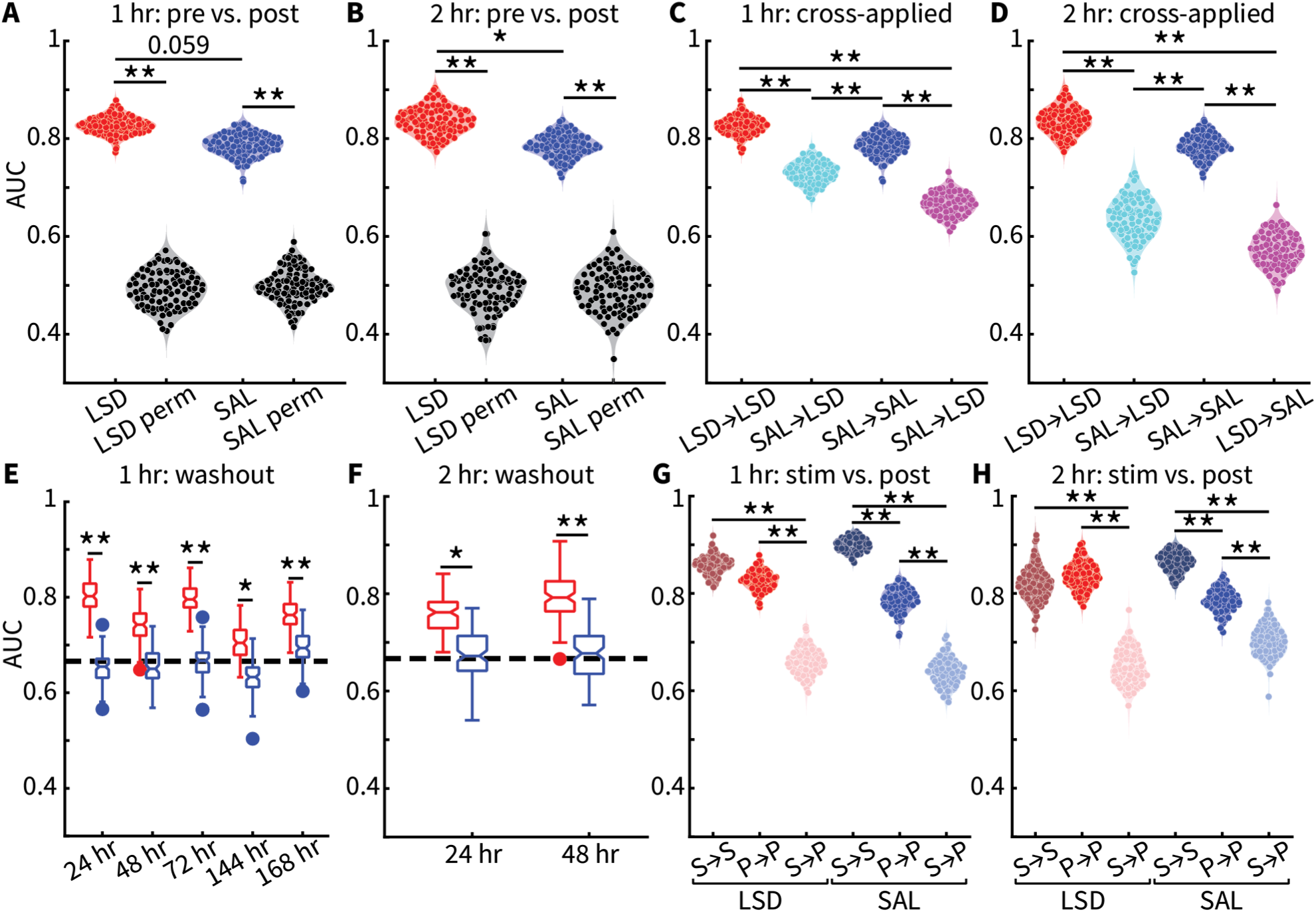
Both cohorts recapitulate the results of the combined cohorts. **A-B.** Pre vs. post models outperform chance (black distributions) and LSD outperforms saline (SAL) at a trend level for cohort 1 (1 hour stimulation; **A**) and significantly for cohort 2 (2 hours stimulation; **B**). **C-D.** Cross-applied models decrease performance in both cohorts. **E-F.** LSD pre vs. post models applied to washout days outperform SAL models and overfit control (heavy dashed line). **G-H.** Pre vs. stim models applied to pre vs. post data (S→P) results in a significant decrease in performance compared to either model tested on its own data (S→S or P→P) in both cohorts. ** p < 0.01, * p < 0.05

