## Supplementary Materials for "Lysergic acid diethylamide pretreatment prolongs brain-stimulation induced neural activity changes"

| Cohort 1: 1 hour stim | sex | DOB | intervention start | age at start | last recording of first intervention | second intervention |
|---|---|---|---|---|---|---|
| LSD25 | F | 2024-02-01 | 2024-06-04 | 124 | 2024-06-11 | 2024-07-22 |
| LSD26 | F | 2024-02-01 | 2024-06-04 | 124 | 2024-06-11 | 2024-07-22 |
| LSD27 | F | 2024-02-01 | 2024-06-04 | 124 | 2024-06-11 | 2024-07-22 |
| LSD28 | F | 2024-02-01 | 2024-06-04 | 124 | 2024-06-11 | 2024-07-22 |
| LSD29 | M | 2024-02-01 | 2024-06-05 | 125 | 2024-06-12 | 2024-07-23 |
| LSD30 | M | 2024-02-01 | 2024-06-05 | 125 | 2024-06-12 | 2024-07-23 |
| LSD31 | M | 2024-02-01 | 2024-06-05 | 125 | 2024-06-12 | 2024-07-23 |
| LSD32 | M | 2024-02-01 | 2024-06-05 | 125 | 2024-06-12 | 2024-07-23 |
| Cohort 2: 2 hour stim | | | | | | |
| IRDM2 | M | 2018-01-20 | 2019-02-12 | 388 | | |
| IRDM21 | F | 2018-07-23 | 2019-06-06 | 318 | | |
| IRDM5 | M | 2018-01-20 | 2019-02-12 | 388 | | |
| IRDM6 | M | 2018-01-20 | 2019-02-12 | 388 | | |
| LSD33 | F | 2024-08-01 | 2024-11-25 | 116 | | |
| LSD34 | F | 2024-08-01 | 2024-11-25 | 116 | | |
| LSD35 | F | 2024-08-01 | 2024-11-25 | 116 | | |
| LSD36 | M | 2024-08-01 | 2024-11-25 | 116 | | |
| LSD37 | M | 2024-08-01 | 2024-11-25 | 116 | | |
| LSD38 | M | 2024-08-01 | 2024-11-25 | 116 | | |
| Cohort 3: 1 hour stim then perfusion | | | | | | |
| LSD09 | M | 2023-04-03 | 2024-01-05 | 277 | SAL-sIL | |
| LSD10 | M | 2023-04-03 | 2024-01-05 | 277 | LSD-sIL | |
| LSD11 | M | 2023-04-03 | 2024-01-05 | 277 | LSD-sham | |
| LSD12 | M | 2023-04-03 | 2024-01-05 | 277 | SAL-sham | |
| LSD13 | F | 2023-04-03 | 2024-01-05 | 277 | SAL-sham | |
| LSD14 | F | 2023-04-03 | 2024-01-05 | 277 | LSD-sIL | |
| LSD15 | F | 2023-04-03 | 2024-01-05 | 277 | SAL-sIL | |

| | | | | | |
|---|---|---|---|---|---|
| LSD17 | F | 2023-10-19 | 2024-03-08 | 141 | SAL-sham |
| LSD18 | F | 2023-10-19 | 2024-03-08 | 141 | SAL-sIL |
| LSD19 | F | 2023-10-19 | 2024-03-14 | 147 | LSD-sham |
| LSD20 | F | 2023-10-19 | 2024-03-14 | 147 | LSD-sIL |
| LSD21 | M | 2023-10-19 | 2024-03-20 | 153 | SAL-sham |
| LSD22 | M | 2023-10-19 | 2024-03-20 | 153 | SAL-sIL |
| LSD23 | M | 2023-10-19 | 2024-03-21 | 154 | LSD-sham |
| IRDM118 | M | 2022-08-11 | 2023-09-05 | 390 | LSD-sham |
| IRDM124 | F | 2022-08-11 | 2023-09-05 | 390 | LSD-sIL |
| IRDM125 | F | 2022-08-11 | 2023-09-05 | 390 | SAL-sIL |
| IRDM136 | F | 2022-12-01 | 2024-04-02 | 488 | LSD-sham |
| IRDM139 | M | 2022-11-24 | 2024-04-02 | 495 | SAL-sIL |
| IRDM140 | M | 2022-11-24 | 2024-04-03 | 496 | SAL-sham |
| IRDM141 | M | 2022-11-24 | 2024-04-02 | 495 | LSD-sham |

| **Figure 2A: pre vs. post stim** | | | |
|---|---|---|---|
| *permutation test* | m+1 | b+1 | p value |
| LSD vs. permuted (not shown) | 1 | 101 | 0.010 |
| SAL vs. permuted (not shown) | 1 | 101 | 0.010 |
| LSD vs. SAL | 1 | 101 | 0.010 |
| *group statistics* | mean | STD | |
| LSD | 0.794 | 0.017 | |
| permuted LSD | 0.504 | 0.030 | |
| SAL | 0.754 | 0.016 | |
| permuted SAL | 0.503 | 0.033 | |

| **Figure 2B: washout days** | | | | | | | |
|---|---|---|---|---|---|---|---|
| *mixed effects model* | | | | | | | |
| *auc~group*time* | estimate | SE | tStat | DF | p value | lower | upper |
| Intercept | 0.642 | 0.028 | 23.157 | 6 | 4.25E-07 | 0.574 | 0.710 |
| group | 0.124 | 0.039 | 3.160 | 6 | 0.020 | 0.028 | 0.220 |
| time | 0.002 | 0.008 | 0.188 | 6 | 0.857 | -0.019 | 0.022 |
| group:time | -0.022 | 0.012 | -1.818 | 6 | 0.119 | -0.050 | 0.007 |
| *permutation tests* | m+1 | b+1 | p value | | | | |
| LSD vs. SAL: 24 hours | 1 | 101 | 0.010 | | | | |
| LSD vs. SAL: 48 hours | 3 | 101 | 0.030 | | | | |
| LSD vs. SAL: 72 hours | 2 | 101 | 0.020 | | | | |
| LSD vs. SAL: 144 hours | 24 | 101 | 0.238 | | | | |
| LSD vs. SAL: 168 hours | 17 | 101 | 0.168 | | | | |
| *group statistics* | | | | | | | |
| 24 hours | mean | STD | | 144 hours | mean | STD | |

| | | | | | |
|---|---|---|---|---|---|
| LSD | 0.770 | 0.024 | LSD | 0.638 | 0.035 |
| LSD perm | 0.492 | 0.050 | LSD perm | 0.495 | 0.052 |
| SAL | 0.647 | 0.025 | SAL | 0.615 | 0.031 |
| SAL perm | 0.501 | 0.045 | SAL perm | 0.499 | 0.048 |
| <u>48 hours</u> | mean | STD | <u>168 hours</u> | mean | STD |
| LSD | 0.697 | 0.033 | LSD | 0.700 | 0.030 |
| LSD perm | 0.502 | 0.049 | LSD perm | 0.495 | 0.056 |
| SAL | 0.641 | 0.031 | SAL | 0.668 | 0.030 |
| SAL perm | 0.506 | 0.053 | SAL perm | 0.500 | 0.053 |
| <u>72 hours</u> | mean | STD | | | |
| LSD | 0.726 | 0.029 | | | |
| LSD perm | 0.495 | 0.059 | | | |
| SAL | 0.663 | 0.031 | | | |
| SAL perm | 0.500 | 0.053 | | | |

| **Figure 2E: variability across time** | | | | | |
|---|---|---|---|---|---|
| *2 sample t-test* | p value | CI | tStat | DF | STD |
| LSD vs. SAL | 0.0343 | [-0.033 -0.001] | -2.1932 | 39 | 0.025 |
| *group statistics* | mean | STD | mean difference | | |
| LSD | 0.050 | 0.132 | -0.017 | | |
| SAL | 0.067 | 0.031 | | | |

| **Figure 2F: variability across rats** | | | | | |
|---|---|---|---|---|---|
| *2 sample t-test* | p value | CI | tStat | DF | STD |
| LSD vs. SAL | 0.00001 | [-0.058 -0.027] | -5.607 | 22 | 0.019 |
| *group statistics* | mean | STD | mean difference | | |

| | | | | |
|---|---|---|---|---|
| | LSD | 0.083 | 0.014 | -0.042 |
| | SAL | 0.125 | 0.022 | |

**Figure 3A-C: stable wash features**

| LSD post-wash | | | SAL post-wash | | |
|---|---|---|---|---|---|
| **176** | **rOFCrNAcCt** | **0.679** | 158 | rOFClNAcSt | 0.623 |
| **33** | **rNAcSa** | **-0.675** | 27 | lNAcSa | -0.623 |
| **27** | **lNAcSa** | **-0.674** | 104 | rmPFClNAcSt | 0.621 |
| **170** | **rOFClNAcCt** | **0.666** | 33 | rNAcSa | -0.620 |
| **9** | **rmPFCa** | **-0.666** | 164 | rOFCrNAcSt | 0.620 |
| **175** | **rOFCrNAcCd** | **0.655** | 24 | rOFChg | 0.618 |
| **102** | **rmPFCrOFChg** | **0.650** | 116 | rmPFClNAcCt | 0.617 |
| **60** | **lmPFClOFChg** | **0.649** | 30 | lNAcShg | 0.617 |
| **10** | **rmPFCb** | **-0.647** | 26 | lNAcSt | -0.616 |
| 164 | rOFCrNAcSt | 0.643 | 170 | rOFClNAcCt | 0.613 |
| **152** | **lOFCrNAcCt** | **0.640** | 68 | lmPFClNAcSt | 0.610 |
| **194** | **lNAcSrNAcCt** | **0.639** | 36 | rNAcShg | 0.607 |
| 158 | rOFClNAcSt | 0.638 | 134 | lOFClNAcSt | 0.605 |
| **206** | **rNAcSrNAcCt** | **0.631** | 74 | lmPFCrNAcSt | 0.602 |
| 169 | rOFClNAcCd | 0.627 | 176 | rOFCrNAcCt | 0.601 |
| 15 | lOFCa | -0.623 | 140 | lOFCrNAcSt | 0.600 |
| 11 | rmPFClg | 0.622 | 21 | rOFCa | -0.598 |
| 205 | rNAcSrNAcCd | 0.622 | 80 | lmPFClNAcCt | 0.597 |
| 200 | rNAcSlNAcCt | 0.622 | 122 | rmPFCrNAcCt | 0.594 |
| 146 | lOFClNAcCt | 0.621 | 9 | rmPFCa | -0.591 |
| 140 | lOFCrNAcSt | 0.613 | 39 | lNAcCa | -0.590 |
| 134 | lOFClNAcSt | 0.612 | 146 | lOFClNAcCt | 0.589 |
| 188 | lNAcSlNAcCt | 0.610 | 206 | rNAcSrNAcCt | 0.586 |
| 193 | lNAcSrNAcCd | 0.607 | 15 | lOFCa | -0.584 |
| 3 | lmPFCa | -0.605 | 152 | lOFCrNAcCt | 0.583 |
| 21 | rOFCa | -0.605 | 38 | lNAcCt | -0.578 |
| 98 | rmPFCrOFCt | -0.604 | 19 | rOFCd | -0.577 |
| 157 | rOFClNAcSd | 0.603 | 45 | rNAcCa | -0.574 |
| 22 | rOFCb | -0.601 | 86 | lmPFCrNAcCt | 0.574 |
| 163 | rOFCrNAcSd | 0.601 | 60 | lmPFClOFChg | 0.569 |
| 92 | rmPFClOFCt | -0.598 | 12 | rmPFChg | 0.569 |
| 45 | rNAcCa | -0.598 | 42 | lNAcChg | 0.569 |
| 122 | rmPFCrNAcCt | 0.595 | 44 | rNAcCt | -0.566 |
| 50 | lmPFCrmPFCt | 0.595 | 32 | rNAcSt | -0.564 |
| 101 | rmPFCrOFClg | 0.594 | 3 | lmPFCa | -0.563 |
| 86 | lmPFCrNAcCt | 0.593 | 149 | lOFClNAcClg | -0.561 |
| 4 | lmPFCb | -0.593 | 28 | lNAcSb | -0.561 |
| 49 | lmPFCrmPFCd | 0.589 | 188 | lNAcSlNAcCt | 0.560 |
| 36 | rNAcShg | 0.587 | 101 | rmPFCrOFClg | 0.560 |
| 116 | rmPFClNAcCt | 0.584 | 98 | rmPFCrOFCt | -0.560 |
| 151 | lOFCrNAcCd | 0.583 | 179 | rOFCrNAcClg | -0.559 |

legend

**bold** = outperforms SAL

shaded = outperforms permuted

l = left; r = right

IL = infralimbic cortex

OFC = orbitofronal cortex

NAcC = nucleus accumbens core

NAcS = nucleus accumbens shell

**Figure 3B: LSD features outperforming SAL and permuted**

| *permutation test* | m+1 | b+1 | p value | |
|---|---|---|---|---|
| rOFCrNAcCt | 1 | 101 | 0.010 | |
| rNAcSa | 2 | 101 | 0.020 | |
| lNAcSa | 3 | 101 | 0.030 | |
| rOFClNAcCt | 2 | 101 | 0.020 | |
| rmPFCa | 1 | 101 | 0.010 | |
| rOFCrNAcCd | 1 | 101 | 0.010 | |
| rmPFCrOFChg | 1 | 101 | 0.010 | |
| lmPFClOFChg | 1 | 101 | 0.010 | |
| rmPFCb | 1 | 101 | 0.010 | |
| rOFCrNAcSt | 18 | 101 | 0.178 | |
| lOFCrNAcCt | 1 | 101 | 0.010 | |
| lNAcSrNAcCt | 1 | 101 | 0.010 | |
| rOFClNAcSt | 26 | 101 | 0.257 | |
| rNAcSrNAcCt | 5 | 101 | 0.050 | |
| *group statistics* | LSD (mean) | LSD (STD) | SAL (mean) | SAL (STD) |
| rOFCrNAcCt | 0.679 | 0.024 | 0.601 | 0.024 |
| rNAcSa | 0.675 | 0.030 | 0.620 | 0.026 |
| lNAcSa | 0.674 | 0.031 | 0.623 | 0.025 |
| rOFClNAcCt | 0.666 | 0.027 | 0.613 | 0.025 |
| rmPFCa | 0.666 | 0.027 | 0.591 | 0.026 |
| rOFCrNAcCd | 0.655 | 0.025 | 0.523 | 0.028 |
| rmPFCrOFChg | 0.650 | 0.023 | 0.473 | 0.089 |
| lmPFClOFChg | 0.649 | 0.027 | 0.569 | 0.040 |
| rmPFCb | 0.647 | 0.024 | 0.554 | 0.024 |
| rOFCrNAcSt | 0.643 | 0.025 | 0.620 | 0.023 |
| lOFCrNAcCt | 0.640 | 0.027 | 0.583 | 0.022 |
| lNAcSrNAcCt | 0.639 | 0.026 | 0.524 | 0.062 |
| rOFClNAcSt | 0.638 | 0.028 | 0.623 | 0.024 |
| rNAcSrNAcCt | 0.631 | 0.024 | 0.586 | 0.025 |

**Figure 3D: cross-applied pre vs. post**

| *rNAcSrNAcCt* | m+1 | b+1 | p value |
|---|---|---|---|
| LSD→LSD vs. SAL →LSD | 1 | 101 | 0.010 |
| LSD→LSD vs. LSD→SAL | 1 | 101 | 0.010 |
| SAL→SAL vs. SAL→LSD | 1 | 101 | 0.010 |
| SAL→SAL vs. LSD→SAL | 1 | 101 | 0.010 |
| *group statistics* | mean | STD | |
| LSD→LSD | 0.794 | 0.017 | |
| SAL→LSD | 0.704 | 0.017 | |

| | | | | | |
|---|---|---|---|---|---|
| 85 | lmPFCrNAcCd | 0.582 | 55 | lmPFClOFCd | 0.559 |
| 1 | lmPFCd | 0.582 | 167 | rOFCrNAcSlg | -0.558 |
| 51 | lmPFCrmPFCa | 0.582 | 200 | rNAcSlNAcCt | 0.555 |
| 12 | rmPFChg | 0.579 | 10 | rmPFCb | -0.554 |
| 78 | lmPFCrNAcShg | 0.579 | 187 | lNAcSlNAcCd | 0.551 |
| 191 | lNAcSlNAcClg | 0.578 | 91 | rmPFClOFCd | 0.548 |
| 30 | lNAcShg | 0.577 | 4 | lmPFCb | -0.545 |
| 39 | lNAcCa | -0.574 | 59 | lmPFClOFClg | 0.544 |
| 18 | lOFChg | 0.572 | 11 | rmPFClg | 0.544 |
| 55 | lmPFClOFCd | 0.572 | 73 | lmPFCrNAcSd | 0.543 |
| 44 | rNAcCt | -0.571 | 150 | lOFClNAcChg | -0.541 |
| 148 | lOFClNAcCb | -0.569 | 49 | lmPFCrmPFCd | 0.541 |
| 16 | lOFCb | -0.568 | 193 | lNAcSrNAcCd | 0.541 |
| 80 | lmPFClNAcCt | 0.567 | 79 | lmPFClNAcCd | 0.540 |
| 171 | rOFClNAcCa | -0.567 | 169 | rOFClNAcCd | 0.540 |
| 38 | lNAcCt | -0.565 | 210 | rNAcSrNAcChg | 0.539 |
| 40 | lNAcCb | 0.564 | 48 | rNAcChg | 0.538 |
| 72 | lmPFClNAcShg | 0.563 | 205 | rNAcSrNAcCd | 0.537 |
| 199 | rNAcSlNAcCd | 0.561 | 22 | rOFCb | -0.534 |
| 67 | lmPFClNAcSd | 0.561 | 163 | rOFCrNAcSd | 0.533 |
| 58 | lmPFClOFCb | 0.561 | 199 | rNAcSlNAcCd | 0.532 |
| 97 | rmPFCrOFCd | -0.561 | 57 | lmPFClOFCa | -0.532 |
| 74 | lmPFCrNAcSt | 0.560 | 62 | lmPFCrOFCt | -0.532 |
| 154 | lOFCrNAcCb | -0.560 | 137 | lOFClNAcSlg | -0.531 |
| 117 | rmPFClNAcCa | -0.560 | 43 | rNAcCd | -0.531 |
| 52 | lmPFCrmPFCb | 0.555 | 85 | lmPFCrNAcCd | 0.531 |
| 167 | rOFCrNAcSlg | -0.550 | 143 | lOFCrNAcSlg | -0.529 |
| 121 | rmPFCrNAcCd | 0.549 | 92 | rmPFClOFCt | -0.528 |
| 84 | lmPFClNAcChg | 0.548 | 67 | lmPFClNAcSd | 0.528 |
| 24 | rOFChg | 0.547 | 56 | lmPFClOFCt | -0.528 |
| 192 | lNAcSlNAcChg | 0.547 | 63 | lmPFCrOFCa | -0.526 |
| 79 | lmPFClNAcCd | 0.547 | 99 | rmPFCrOFCa | -0.526 |
| 111 | rmPFCrNAcSa | -0.545 | 148 | lOFClNAcCb | -0.524 |
| 131 | lOFCrOFClg | 0.545 | 194 | lNAcSrNAcCt | 0.524 |
| 65 | lmPFCrOFClg | 0.544 | 121 | rmPFCrNAcCd | 0.523 |
| 196 | lNAcSrNAcCb | -0.543 | 61 | lmPFCrOFCd | 0.523 |
| 153 | lOFCrNAcCa | -0.542 | 175 | rOFCrNAcCd | 0.523 |
| 145 | lOFClNAcCd | 0.542 | 115 | rmPFClNAcCd | 0.522 |
| 187 | lNAcSlNAcCd | 0.542 | 157 | rOFClNAcSd | 0.522 |
| 118 | rmPFClNAcCb | -0.541 | 165 | rOFCrNAcSa | 0.521 |
| 161 | rOFClNAcSlg | -0.541 | 46 | rNAcCb | 0.520 |
| 59 | lmPFClOFClg | 0.540 | 83 | lmPFClNAcClg | -0.519 |
| 123 | rmPFCrNAcCa | -0.540 | 204 | rNAcSlNAcChg | -0.519 |
| 57 | lmPFClOFCa | 0.540 | 90 | lmPFCrNAcChg | 0.519 |

| | | |
|---|---|---|
| LSD→SAL | 0.647 | 0.020 |
| SAL→SAL | 0.754 | 0.016 |

| **Figure 3E: LSD vs. SAL** | | | |
|---|---|---|---|
| *permutation test* | m+1 | b+1 | p value |
| stim vs. permuted | 1 | 101 | 0.010 |
| post vs. permuted | 1 | 101 | 0.010 |
| *group statistics* | mean | STD | |
| stim | 0.800 | 0.016 | |
| stim permuted | 0.499 | 0.031 | |
| post | 0.752 | 0.020 | |
| post permuted | 0.500 | 0.030 | |

| | | | | | |
|---|---|---|---|---|---|
| 105 | rmPFClNAcSa | -0.540 | 37 | lNAcCd | -0.519 |
| 136 | lOFClNAcSb | -0.539 | 154 | lOFCrNAcCb | -0.519 |
| 68 | lmPFClNAcSt | 0.538 | 192 | lNAcSlNAcChg | 0.519 |
| 177 | rOFCrNAcCa | -0.538 | 133 | lOFClNAcSd | -0.517 |
| 62 | lmPFCrOFCt | -0.535 | 82 | lmPFClNAcCb | 0.517 |
| 159 | rOFClNAcSa | -0.535 | 198 | lNAcSrNAcChg | -0.517 |
| 178 | rOFCrNAcCb | -0.534 | 135 | lOFClNAcSa | 0.517 |
| 63 | lmPFCrOFCa | 0.533 | 215 | lNAcCrNAcClg | -0.516 |
| 172 | rOFClNAcCb | -0.532 | 197 | lNAcSrNAcClg | -0.516 |
| 66 | lmPFCrOFChg | 0.532 | 214 | lNAcCrNAcCb | -0.516 |
| 119 | rmPFClNAcClg | 0.531 | 129 | lOFCrOFCa | -0.515 |
| 2 | lmPFCt | 0.530 | 81 | lmPFClNAcCa | -0.515 |
| 202 | rNAcSlNAcCb | -0.529 | 151 | lOFCrNAcCd | 0.515 |
| 147 | lOFClNAcCa | -0.528 | 65 | lmPFCrOFClg | 0.515 |
| 90 | lmPFCrNAcChg | 0.527 | 153 | lOFCrNAcCa | -0.514 |
| 46 | rNAcCb | 0.527 | 77 | lmPFCrNAcSlg | -0.513 |
| 203 | rNAcSlNAcClg | 0.526 | 78 | lmPFCrNAcShg | 0.511 |
| 174 | rOFClNAcChg | 0.526 | 185 | lNAcSrNAcSlg | -0.511 |
| 124 | rmPFCrNAcCb | -0.526 | 125 | rmPFCrNAcClg | -0.511 |
| 99 | rmPFCrOFCa | -0.525 | 139 | lOFCrNAcSd | -0.510 |
| 42 | lNAcChg | -0.523 | 93 | rmPFClOFCa | -0.509 |
| 94 | rmPFClOFCb | 0.522 | 145 | lOFClNAcCd | 0.509 |
| 56 | lmPFClOFCt | -0.522 | 94 | rmPFClOFCb | 0.509 |
| 204 | rNAcSlNAcChg | -0.522 | 160 | rOFClNAcSb | -0.509 |
| 106 | rmPFClNAcSb | -0.521 | 114 | rmPFCrNAcShg | -0.509 |
| 142 | lOFCrNAcSb | -0.521 | 127 | lOFCrOFCd | 0.000 |
| 115 | rmPFClNAcCd | 0.521 | 142 | lOFCrNAcSb | -0.508 |
| 189 | lNAcSlNAcCa | -0.521 | 95 | rmPFClOFClg | 0.507 |
| 89 | lmPFCrNAcClg | 0.519 | 196 | lNAcSrNAcCb | -0.506 |
| 7 | rmPFCd | 0.519 | 87 | lmPFCrNAcCa | -0.506 |
| 190 | lNAcSlNAcCb | -0.519 | 130 | lOFCrOFCb | 0.505 |
| 181 | lNAcSrNAcSd | -0.519 | 70 | lmPFClNAcSb | 0.505 |
| 77 | lmPFCrNAcSlg | 0.517 | 50 | lmPFCrmPFCt | 0.505 |
| 201 | rNAcSlNAcCa | -0.517 | 53 | lmPFCrmPFClg | -0.504 |
| 179 | rOFCrNAcClg | -0.516 | 161 | rOFClNAcSlg | -0.504 |
| 6 | lmPFChg | -0.516 | 131 | lOFCrOFClg | 0.503 |
| 155 | lOFCrNAcClg | 0.515 | 64 | lmPFCrOFCb | 0.503 |
| 64 | lmPFCrOFCb | 0.515 | 51 | lmPFCrmPFCa | -0.503 |
| 165 | rOFCrNAcSa | -0.513 | 159 | rOFClNAcSa | 0.503 |
| 141 | lOFCrNAcSa | -0.512 | 209 | rNAcSrNAcClg | 0.502 |
| 93 | rmPFClOFCa | -0.512 | 156 | lOFCrNAcChg | 0.502 |
| 91 | rmPFClOFCd | 0.512 | 203 | rNAcSlNAcClg | 0.502 |
| 83 | lmPFClNAcClg | 0.512 | 177 | rOFCrNAcCa | 0.501 |
| 81 | lmPFClNAcCa | -0.512 | 191 | lNAcSlNAcClg | 0.501 |

| | | | | | |
|---|---|---|---|---|---|
| 130 | lOFCrOFCb | -0.511 | 47 | rNAcClg | 0.501 |
| 13 | lOFCd | 0.511 | 126 | rmPFCrNAcChg | -0.500 |
| 209 | rNAcSrNAcClg | 0.511 | 97 | rmPFCrOFCd | -0.500 |
| 31 | rNAcSd | -0.510 | 100 | rmPFCrOFCb | -0.500 |
| 208 | rNAcSrNAcCb | -0.510 | 213 | lNAcCrNAcCa | -0.500 |
| 73 | lmPFCrNAcSd | 0.508 | 171 | rOFClNAcCa | -0.500 |
| 135 | lOFClNAcSa | -0.507 | 54 | lmPFCrmPFChg | 0.499 |
| 82 | lmPFClNAcCb | 0.506 | 147 | lOFClNAcCa | 0.498 |
| 88 | lmPFCrNAcCb | -0.506 | 117 | rmPFClNAcCa | -0.498 |
| 53 | lmPFCrmPFClg | 0.505 | 120 | rmPFClNAcChg | 0.000 |
| 61 | lmPFCrOFCd | 0.505 | 136 | lOFClNAcSb | -0.498 |
| 112 | rmPFCrNAcSb | -0.505 | 34 | rNAcSb | -0.497 |
| 54 | lmPFCrmPFChg | 0.504 | 72 | lmPFClNAcShg | -0.497 |
| 70 | lmPFClNAcSb | 0.504 | 172 | rOFClNAcCb | 0.497 |
| 113 | rmPFCrNAcSlg | 0.503 | 112 | rmPFCrNAcSb | 0.497 |
| 109 | rmPFCrNAcSd | -0.503 | 201 | rNAcSlNAcCa | -0.497 |
| 160 | rOFClNAcSb | -0.502 | 75 | lmPFCrNAcSa | -0.497 |
| 104 | rmPFClNAcSt | -0.502 | 69 | lmPFClNAcSa | -0.497 |
| 114 | rmPFCrNAcShg | 0.502 | 84 | lmPFClNAcChg | -0.497 |
| 76 | lmPFCrNAcSb | 0.502 | 141 | lOFCrNAcSa | -0.497 |
| 156 | lOFCrNAcChg | 0.501 | 182 | lNAcSrNAcSt | -0.497 |
| 107 | rmPFClNAcSlg | -0.501 | 103 | rmPFClNAcSd | -0.497 |
| 183 | lNAcSrNAcSa | -0.500 | 174 | rOFClNAcChg | 0.496 |
| 133 | lOFClNAcSd | -0.499 | 211 | lNAcCrNAcCd | -0.496 |
| 48 | rNAcChg | -0.499 | 40 | lNAcCb | 0.496 |
| 71 | lmPFClNAcSlg | 0.499 | 216 | lNAcCrNAcChg | -0.495 |
| 195 | lNAcSrNAcCa | -0.498 | 184 | lNAcSrNAcSb | -0.495 |
| 95 | rmPFClOFClg | 0.498 | 207 | rNAcSrNAcCa | -0.494 |
| 214 | lNAcCrNAcCb | -0.498 | 132 | lOFCrOFChg | -0.492 |
| 32 | rNAcSt | 0.498 | 183 | lNAcSrNAcSa | 0.000 |
| 186 | lNAcSrNAcShg | 0.498 | 123 | rmPFCrNAcCa | -0.492 |
| 184 | lNAcSrNAcSb | -0.497 | 195 | lNAcSrNAcCa | -0.491 |
| 100 | rmPFCrOFCb | -0.497 | 189 | lNAcSlNAcCa | -0.491 |
| 143 | lOFCrNAcSlg | 0.496 | 52 | lmPFCrmPFCb | 0.491 |
| 216 | lNAcCrNAcChg | 0.496 | 155 | lOFCrNAcClg | 0.488 |
| 87 | lmPFCrNAcCa | -0.496 | 212 | lNAcCrNAcCt | -0.487 |
| 19 | rOFCd | 0.495 | 108 | rmPFClNAcShg | -0.487 |
| 69 | lmPFClNAcSa | -0.495 | 166 | rOFCrNAcSb | 0.487 |
| 129 | lOFCrOFCa | -0.495 | 58 | lmPFClOFCb | 0.487 |
| 139 | lOFCrNAcSd | -0.494 | 181 | lNAcSrNAcSd | -0.487 |
| 137 | lOFClNAcSlg | 0.494 | 31 | rNAcSd | 0.487 |
| 185 | lNAcSrNAcSlg | 0.494 | 144 | lOFCrNAcShg | 0.487 |
| 108 | rmPFClNAcShg | -0.494 | 109 | rmPFCrNAcSd | -0.486 |
| 128 | lOFCrOFCt | 0.493 | 23 | rOFClg | -0.486 |

| | | | | | |
|---|---|---|---|---|---|
| 215 | lNAcCrNAcClg | -0.492 | 17 | lOFClg | -0.485 |
| 25 | lNAcSd | 0.492 | 186 | lNAcSrNAcShg | 0.484 |
| 120 | rmPFClNAcChg | -0.491 | 96 | rmPFClOFChg | -0.484 |
| 132 | lOFCrOFChg | -0.490 | 76 | lmPFCrNAcSb | -0.484 |
| 207 | rNAcSrNAcCa | -0.489 | 113 | rmPFCrNAcSlg | 0.483 |
| 125 | rmPFCrNAcClg | 0.489 | 1 | lmPFCd | 0.482 |
| 173 | rOFClNAcClg | 0.487 | 202 | rNAcSlNAcCb | -0.482 |
| 213 | lNAcCrNAcCa | -0.486 | 71 | lmPFClNAcSlg | 0.482 |
| 103 | rmPFClNAcSd | -0.485 | 16 | lOFCb | 0.482 |
| 162 | rOFClNAcShg | 0.484 | 107 | rmPFClNAcSlg | 0.480 |
| 210 | rNAcSrNAcChg | 0.484 | 128 | lOFCrOFCt | 0.480 |
| 212 | lNAcCrNAcCt | -0.483 | 88 | lmPFCrNAcCb | -0.480 |
| 166 | rOFCrNAcSb | -0.480 | 89 | lmPFCrNAcClg | 0.477 |
| 144 | lOFCrNAcShg | 0.478 | 105 | rmPFClNAcSa | -0.476 |
| 198 | lNAcSrNAcChg | 0.478 | 41 | lNAcClg | -0.476 |
| 75 | lmPFCrNAcSa | -0.478 | 208 | rNAcSrNAcCb | -0.475 |
| 197 | lNAcSrNAcClg | 0.477 | 119 | rmPFClNAcClg | 0.474 |
| 37 | lNAcCd | -0.475 | 124 | rmPFCrNAcCb | -0.474 |
| 20 | rOFCt | 0.471 | 102 | rmPFCrOFChg | -0.473 |
| 43 | rNAcCd | -0.470 | 111 | rmPFCrNAcSa | -0.472 |
| 182 | lNAcSrNAcSt | -0.470 | 106 | rmPFClNAcSb | -0.471 |
| 126 | rmPFCrNAcChg | 0.465 | 5 | lmPFClg | -0.470 |
| 14 | lOFCt | 0.460 | 25 | lNAcSd | 0.468 |
| 110 | rmPFCrNAcSt | -0.457 | 66 | lmPFCrOFChg | -0.468 |
| 149 | lOFClNAcClg | 0.457 | 162 | rOFClNAcShg | 0.462 |
| 168 | rOFCrNAcShg | 0.453 | 178 | rOFCrNAcCb | 0.462 |
| 180 | rOFCrNAcChg | 0.452 | 118 | rmPFClNAcCb | -0.461 |
| 127 | lOFCrOFCd | 0.452 | 35 | rNAcSlg | -0.460 |
| 96 | rmPFClOFChg | -0.449 | 173 | rOFClNAcClg | 0.460 |
| 8 | rmPFCt | 0.439 | 138 | lOFClNAcShg | 0.459 |
| 17 | lOFClg | -0.437 | 2 | lmPFCt | 0.457 |
| 41 | lNAcClg | -0.437 | 7 | rmPFCd | 0.452 |
| 26 | lNAcSt | 0.437 | 190 | lNAcSlNAcCb | -0.449 |
| 47 | rNAcClg | -0.435 | 6 | lmPFChg | -0.446 |
| 5 | lmPFClg | -0.434 | 29 | lNAcSlg | -0.445 |
| 138 | lOFClNAcShg | 0.431 | 8 | rmPFCt | 0.442 |
| 35 | rNAcSlg | -0.427 | 180 | rOFCrNAcChg | 0.439 |
| 23 | rOFClg | -0.411 | 168 | rOFCrNAcShg | 0.438 |
| 211 | lNAcCrNAcCd | -0.410 | 110 | rmPFCrNAcSt | -0.434 |
| 34 | rNAcSb | 0.392 | 13 | lOFCd | 0.428 |
| 29 | lNAcSlg | -0.385 | 20 | rOFCt | 0.428 |
| 150 | lOFClNAcChg | 0.368 | 14 | lOFCt | 0.426 |
| 28 | lNAcSb | 0.365 | 18 | lOFChg | -0.401 |

**Figure 4A**

| | | | |
|---|---|---|---|
| *permutation test* | | | |
| LSD | m+1 | b+1 | p |
| stim→stim vs. post→post | 1 | 101 | 0.010 |
| stim→stim vs. stim→post | 1 | 101 | 0.010 |
| post→post vs. stim→post | 1 | 101 | 0.010 |
| SAL | m+1 | b+1 | p |
| stim→stim vs. post→post | 1 | 101 | 0.010 |
| stim→stim vs. stim→post | 1 | 101 | 0.010 |
| post→post vs. stim→post | 1 | 101 | 0.010 |
| *group statistics* | | | |
| LSD | mean | STD | |
| stim→stim | 0.863 | 0.013 | |
| stim→post | 0.661 | 0.022 | |
| post→post | 0.794 | 0.017 | |
| SAL | mean | STD | |
| stim→stim | 0.861 | 0.013 | |
| stim→post | 0.653 | 0.019 | |
| post→post | 0.754 | 0.016 | |

**Figure 4B**

LSD post→wash vs. stim→wash

*mixed effects model*

| *auc~group*time* | estimate | SE | tStat | DF | p value | lower | upper |
|---|---|---|---|---|---|---|---|
| Intercept | 0.766 | 0.026 | 29.361 | 6 | 1.03E-07 | 0.702 | 0.830 |
| group | -0.133 | 0.037 | -3.595 | 6 | 0.011 | -0.223 | -0.042 |

| | | | | | | | |
|---|---|---|---|---|---|---|---|
| time | -0.020 | 0.008 | -2.533 | 6 | 0.045 | -0.039 | -6.77E-04 |
| group:time | 0.006 | 0.011 | 0.528 | 6 | 0.617 | -0.021 | 0.033 |
| <u>SAL post→wash vs. stim→wash</u> | | | | | | | |
| *mixed effects model* | | | | | | | |
| *auc~group*time* | estimate | SE | tStat | DF | p value | lower | upper |
| Intercept | 0.642 | 0.020 | 32.538 | 6 | 5.60E-08 | 0.594 | 0.690 |
| group | -0.015 | 0.028 | -0.548 | 6 | 0.604 | -0.084 | 0.053 |
| time | 0.002 | 0.006 | 0.264 | 6 | 0.801 | -0.013 | 0.016 |
| group:time | -0.008 | 0.008 | -0.921 | 6 | 0.392 | -0.028 | 0.013 |

| Figure 5A: pS6 -- IL | | | | |
|---|---|---|---|---|
| *GLME (Mean ~ Sex + Age + Group); distribution = gamma; link = log* | | | | |
| | FStat | DF1 | DF2 | p value |
| intercept | 3856.5 | 1 | 1156 | 0 |
| group | 4.319 | 3 | 1156 | 0.005 |
| age (z-score) | 9.026 | 1 | 1156 | 0.003 |
| sex | 2.163 | 1 | 1156 | 0.142 |
| Post hoc comparisons | | | | |
| *coefTest* | p value | | | |
| LSD-sIL vs. LSD-sham | 0.001 | | | |
| SAL-sIL vs. LSD-sham | 0.011 | | | |
| SAL-sham vs. LSD-sham | 0.173 | | | |
| SAL-sIL vs. LSD-sIL | 0.307 | | | |
| SAL-sham vs. LSD-sIL | 0.036 | | | |
| SAL-sham vs. SAL-sIL | 0.252 | | | |

| Figure 5B: pS6 -- IL | | | | |
|---|---|---|---|---|
| *GLME (Mean ~ Sex + Age + Group); distribution = gamma; link = log* | | | | |
| | FStat | DF1 | DF2 | p value |
| intercept | 6163.7 | 1 | 749 | 0 |
| group | 6.403 | 3 | 749 | 2.76E-04 |
| age (z-score) | 35.694 | 1 | 749 | 3.57E-09 |
| sex | 9.935 | 1 | 749 | 0.002 |
| Post hoc comparisons | | | | |
| *coefTest* | p value | | | |
| LSD-sIL vs. LSD-sham | 2.34E-05 | | | |
| SAL-sIL vs. LSD-sham | 0.002 | | | |
| SAL-sham vs. LSD-sham | 0.026 | | | |
| SAL-sIL vs. LSD-sIL | 0.262 | | | |
| SAL-sham vs. LSD-sIL | 0.011 | | | |
| SAL-sham vs. SAL-sIL | 0.186 | | | |

| Figure 5C: pS6 -- DG | | | | |
|---|---|---|---|---|
| *GLME (Mean ~ Sex + Age + Group); distribution = gamma; link = log* | | | | |
| | FStat | DF1 | DF2 | p value |
| intercept | 2698 | 1 | 260 | 2.70E-139 |

| Figure 5D: PNN -- IL | | | | |
|---|---|---|---|---|
| *GLME (Mean ~ Sex + Age + Group); distribution = gamma; link = log* | | | | |
| | FStat | DF1 | DF2 | p value |
| intercept | 23011 | 1 | 792 | 0 |
| group | 9.968 | 3 | 792 | 1.87E-06 |
| age (z-score) | 0.166 | 1 | 792 | 0.684 |
| sex | 8.312 | 1 | 792 | 0.004 |
| Post hoc comparisons | | | | |
| *coefTest* | p value | | | |
| LSD-sIL vs. LSD-sham | 0.009 | | | |
| SAL-sIL vs. LSD-sham | 1.39E-05 | | | |
| SAL-sham vs. LSD-sham | 5.75E-07 | | | |
| SAL-sIL vs. LSD-sIL | 0.036 | | | |
| SAL-sham vs. LSD-sIL | 0.008 | | | |
| SAL-sham vs. SAL-sIL | 0.479 | | | |

| Figure 5E: PNN -- PL | | | | |
|---|---|---|---|---|
| *GLME (Mean ~ Sex + Age + Group); distribution = gamma; link = log* | | | | |
| | FStat | DF1 | DF2 | p value |
| intercept | 17134 | 1 | 784 | 0 |
| group | 1.2872 | 3 | 784 | 0.278 |
| age (z-score) | 13.598 | 1 | 784 | 2.42E-04 |
| sex | 6.630 | 1 | 784 | 0.010 |
| Post hoc comparisons | | | | |
| *coefTest* | p value | | | |
| LSD-sIL vs. LSD-sham | 0.432 | | | |
| SAL-sIL vs. LSD-sham | 0.092 | | | |
| SAL-sham vs. LSD-sham | 0.737 | | | |
| SAL-sIL vs. LSD-sIL | 0.304 | | | |
| SAL-sham vs. LSD-sIL | 0.533 | | | |
| SAL-sham vs. SAL-sIL | 0.077 | | | |

| Figure 5F: PNN -- DG | | | | |
|---|---|---|---|---|
| *GLME (Mean ~ Sex + Age + Group); distribution = gamma; link = log* | | | | |
| | FStat | DF1 | DF2 | p value |
| intercept | 32238 | 1 | 1519 | 0 |

| | | | | |
|---|---|---|---|---|
| group | 1.422 | 3 | 260 | 0.237 |
| age (z-score) | 19.011 | 1 | 260 | 1.87E-05 |
| sex | 0.583 | 1 | 260 | 0.446 |
| Post hoc comparisons | | | | |
| *coefTest* | p value | | | |
| LSD-sIL vs. LSD-sham | 0.142 | | | |
| SAL-sIL vs. LSD-sham | 0.941 | | | |
| SAL-sham vs. LSD-sham | 0.455 | | | |
| SAL-sIL vs. LSD-sIL | 0.075 | | | |
| SAL-sham vs. LSD-sIL | 0.461 | | | |
| SAL-sham vs. SAL-sIL | 0.295 | | | |

| | | | | |
|---|---|---|---|---|
| group | 8.971 | 3 | 1519 | 6.85E-06 |
| age (z-score) | 5.461 | 1 | 1519 | 0.02 |
| sex | 7.192 | 1 | 1519 | 0.007 |
| Post hoc comparisons | | | | |
| *coefTest* | p value | | | |
| LSD-sIL vs. LSD-sham | 0.029 | | | |
| SAL-sIL vs. LSD-sham | 0.889 | | | |
| SAL-sham vs. LSD-sham | 0.099 | | | |
| SAL-sIL vs. LSD-sIL | 3.45E-06 | | | |
| SAL-sham vs. LSD-sIL | 0.192 | | | |
| SAL-sham vs. SAL-sIL | 5.12E-05 | | | |

| **Extended Data Figure 1** | | |
|---|---|---|
| *group statistics* | mean | STD |
| base | 0.717 | 0.022 |
| 24 hr | 0.665 | 0.023 |
| 48 hr | 0.709 | 0.035 |
| 96 hr | 0.625 | 0.033 |

| | | | |
|---|---|---|---|
| **Extended Data Figure 2** | | | |
| *permutation test* | m+1 | b+1 | p value |
| LSD vs. LSD M | 90 | 101 | 0.891 |
| LSD vs. LSD F | 68 | 101 | 0.673 |
| LSD M vs. LSD F | 25 | 101 | 0.248 |
| SAL vs. SAL M | 35 | 101 | 0.347 |
| SAL vs. SAL F | 6 | 101 | 0.059 |
| SAL M vs. SAL F | 14 | 101 | 0.139 |
| *group statistics* | mean | STD | |
| LSD | 0.794 | 0.017 | |
| LSD M | 0.822 | 0.022 | |
| LSD F | 0.805 | 0.022 | |
| SAL | 0.754 | 0.016 | |
| SAL M | 0.763 | 0.024 | |
| SAL F | 0.786 | 0.021 | |

| **Extended Data Figure 3A** | | |
|---|---|---|
| *group statistics* | mean | STD |
| pre | 9.350 | 3.276 |
| post | 6.781 | 3.396 |
| 24 hr | 8.763 | 3.360 |
| 48 hr | 8.422 | 2.994 |
| 96 hr | 8.421 | 3.434 |

| **Extended Data Figure 3B** | | |
|---|---|---|
| *group statistics* | mean | STD |
| pre | 8.693 | 3.646 |
| post | 5.304 | 3.623 |
| 24 hr | 7.481 | 3.658 |
| 48 hr | 7.463 | 3.120 |
| 144 hr | 7.490 | 3.140 |
| 168 hr | 7.406 | 3.347 |

| **Extended Data Figure 3C** | | |
|---|---|---|
| *group statistics* | mean | STD |
| pre | 9.561 | 3.754 |
| post | 6.045 | 4.204 |
| 24 hr | 7.398 | 4.244 |
| 48 hr | 7.453 | 3.973 |
| 144 hr | 9.449 | 4.529 |
| 168 hr | 8.650 | 4.913 |

| **Extended Data Figure 3D** | | |
|---|---|---|
| *group statistics* | mean | STD |
| pre vs. post | 0.501 | 0.043 |
| 24 hr | 0.482 | 0.059 |
| 48 hr | 0.490 | 0.055 |
| 96 hr | 0.494 | 0.056 |
| permuted | 0.497 | 0.043 |

| **Extended Data Figure 3E** | | |
|---|---|---|
| *group statistics* | mean | STD |
| pre vs. post | 0.543 | 0.037 |
| 24 hr | 0.510 | 0.045 |
| 48 hr | 0.514 | 0.042 |
| 72 hr | 0.510 | 0.044 |
| 144 hr | 0.518 | 0.039 |
| 168 hr | 0.518 | 0.041 |
| permuted | 0.507 | 0.056 |

| **Extended Data Figure 3F** | | |
|---|---|---|
| *group statistics* | mean | STD |
| pre vs. post | 0.520 | 0.037 |
| 24 hr | 0.511 | 0.054 |
| 48 hr | 0.501 | 0.051 |
| 72 hr | 0.493 | 0.051 |
| 144 hr | 0.481 | 0.058 |
| 168 hr | 0.497 | 0.051 |
| permuted | 0.506 | 0.041 |

| **Extended Data Figure 4A: pre vs. post -- 1 hr** | | | |
|---|---|---|---|
| *permutation test* | m+1 | b+1 | p value |
| LSD vs. SAL | 6 | 101 | 0.059 |
| LSD vs. permuted | 1 | 101 | 0.010 |
| SAL vs. permuted | 1 | 101 | 0.010 |
| *group statistics* | mean | STD | |
| LSD | 0.827 | 0.019 | |
| LSD permuted | 0.493 | 0.036 | |
| SAL | 0.787 | 0.023 | |
| SAL permuted | 0.498 | 0.034 | |

| **Extended Data Figure 4B: pre vs. post -- 2 hr** | | | |
|---|---|---|---|
| *permutation test* | m+1 | b+1 | p value |
| LSD vs. SAL | 2 | 101 | 0.020 |
| LSD vs. permuted | 1 | 101 | 0.010 |
| SAL vs. permuted | 1 | 101 | 0.010 |
| *group statistics* | mean | STD | |
| LSD | 0.836 | 0.028 | |
| LSD permuted | 0.490 | 0.044 | |
| SAL | 0.785 | 0.024 | |
| SAL permuted | 0.491 | 0.044 | |

| **Extended Data Figure 4C: cross-applied -- 1 hr** | | | |
|---|---|---|---|
| *permutation test* | m+1 | b+1 | p value |
| LSD→LSD vs. SAL →LSD | 1 | 101 | 0.010 |
| LSD→LSD vs. LSD→SAL | 1 | 101 | 0.010 |
| SAL→SAL vs. SAL→LSD | 1 | 101 | 0.010 |
| SAL→SAL vs. LSD→SAL | 1 | 101 | 0.010 |
| *group statistics* | Mean | STD | |
| LSD→LSD | 0.827 | 0.019 | |
| SAL→LSD | 0.731 | 0.022 | |
| LSD→SAL | 0.667 | 0.023 | |
| SAL→SAL | 0.787 | 0.023 | |

| **Extended Data Figure 4D: cross-applied -- 2 hr** | | | |
|---|---|---|---|
| *permutation test* | m+1 | b+1 | p value |
| LSD→LSD vs. SAL →LSD | 1 | 101 | 0.010 |
| LSD→LSD vs. LSD→SAL | 1 | 101 | 0.010 |
| SAL→SAL vs. SAL→LSD | 1 | 101 | 0.010 |
| SAL→SAL vs. LSD→SAL | 1 | 101 | 0.010 |
| *group statistics* | Mean | STD | |
| LSD→LSD | 0.836 | 0.028 | |
| SAL→LSD | 0.637 | 0.042 | |
| LSD→SAL | 0.572 | 0.034 | |
| SAL→SAL | 0.785 | 0.024 | |

| **Extended Data Figure 4E: washout -- 1 hr** | | | |
|---|---|---|---|
| *permutation tests* | m+1 | b+1 | p value |
| LSD vs. SAL: 24 hours | 1 | 101 | 0.010 |
| LSD vs. SAL: 48 hours | 1 | 101 | 0.010 |
| LSD vs. SAL: 72 hours | 1 | 101 | 0.010 |
| LSD vs. SAL: 144 hours | 5 | 101 | 0.050 |

| **Extended Data Figure 4E: washout -- 2 hr** | | | |
|---|---|---|---|
| *permutation tests* | m+1 | b+1 | p value |
| LSD vs. SAL: 24 hours | 2 | 101 | 0.020 |
| LSD vs. SAL: 48 hours | 1 | 101 | 0.010 |
| *group statistics* | | | |

| | | | |
|---|---|---|---|
| LSD vs. SAL: 168 hours | 4 | 101 | 0.040 |
| *group statistics* | | | |
| 24 hours | mean | STD | |
| LSD | 0.802 | 0.037 | |
| SAL | 0.651 | 0.032 | |
| 48 hours | mean | STD | |
| LSD | 0.742 | 0.034 | |
| SAL | 0.655 | 0.034 | |
| 72 hours | mean | STD | |
| LSD | 0.797 | 0.027 | |
| SAL | 0.664 | 0.035 | |
| 144 hours | mean | STD | |
| LSD | 0.706 | 0.034 | |
| SAL | 0.632 | 0.037 | |
| 168 hours | mean | STD | |
| LSD | 0.762 | 0.030 | |
| SAL | 0.695 | 0.033 | |

| | | |
|---|---|---|
| 24 hours | mean | STD |
| LSD | 0.757 | 0.037 |
| SAL | 0.676 | 0.046 |
| 48 hours | mean | STD |
| LSD | 0.793 | 0.046 |
| SAL | 0.676 | 0.052 |

| **Extended Data Figure 4G: stim vs. post -- 1hr** | | | |
|---|---|---|---|
| *permutation test* | | | |
| LSD | m+1 | b+1 | p |
| stim→stim vs. post→post | 7 | 101 | 0.069 |
| stim→stim vs. stim→post | 1 | 101 | 0.010 |
| post→post vs. stim→post | 1 | 101 | 0.010 |
| SAL | m+1 | b+1 | p |
| stim→stim vs. post→post | 1 | 101 | 0.010 |

| **Extended Data Figure 4H: stim vs. post -- 2hr** | | | |
|---|---|---|---|
| *permutation test* | | | |
| LSD | m+1 | b+1 | p |
| stim→stim vs. post→post | 73 | 101 | 0.723 |
| stim→stim vs. stim→post | 1 | 101 | 0.010 |
| post→post vs. stim→post | 1 | 101 | 0.010 |
| SAL | m+1 | b+1 | p |
| stim→stim vs. post→post | 1 | 101 | 0.010 |

| | | | |
|---|---|---|---|
| stim→stim vs. stim→post | 1 | 101 | 0.010 |
| post→post vs. stim→post | 1 | 101 | 0.010 |
| *group statistics* | | | |
| LSD | mean | STD | |
| stim→stim | 0.858 | 0.019 | |
| stim→post | 0.660 | 0.028 | |
| post→post | 0.827 | 0.019 | |
| SAL | mean | STD | |
| stim→stim | 0.899 | 0.013 | |
| stim→post | 0.638 | 0.027 | |
| post→post | 0.787 | 0.023 | |

| | | | |
|---|---|---|---|
| stim→stim vs. stim→post | 1 | 101 | 0.010 |
| post→post vs. stim→post | 1 | 101 | 0.010 |
| *group statistics* | | | |
| LSD | mean | STD | |
| stim→stim | 0.820 | 0.035 | |
| stim→post | 0.649 | 0.037 | |
| post→post | 0.836 | 0.028 | |
| SAL | mean | STD | |
| stim→stim | 0.864 | 0.020 | |
| stim→post | 0.702 | 0.031 | |
| post→post | 0.785 | 0.024 | |
